# Mechanistic Link between Executive Functions and Higher Cognition: Evidence from Psychometric Modeling and Brain Stimulation

**DOI:** 10.1101/2025.11.03.686235

**Authors:** Marko Živanović, Jovana Bjekić, Saša R. Filipović

**Author notes:** Corresponding author: Dr. Marko Živanović Institute of Psychology & LIRA lab, Department of Psychology, Faculty of Philosophy, University of Belgrade, Belgrade, Serbia, Čika Ljubina 18-20, 11000 Belgrade, Serbia. Author contributions (CRediT)*: MŽ – Conceptualization, Data curation, Formal analysis, Investigation, Methodology, Writing – original draft; JB – Investigation, Methodology, Funding acquisition, Writing – review and editing; SRF – Methodology, Funding acquisition, Writing – review and editing.

## Abstract

Human cognitive abilities depend on flexible coordination across distributed cognitive control systems within the frontoparietal network, yet the causal architecture linking executive functions (EFs) to higher cognition remains debated. We combined psychometric modeling and neuromodulation to test causal contributions of EFs to distinct cognitive abilities. In a psychometric study, we modeled Unity–Diversity EF components (updating, inhibition, shifting) against four Cattell–Horn–Carroll (CHC) group factors. Fluid reasoning (*Gf*) was uniquely related to both Common EF and Updating-specific variance, whereas visual processing (*Gv*) and processing speed (*Gs*) were primarily linked to Common EF; crystallized ability (*Gc*) proved to be largely independent of EFs. To causally probe this architecture, in neuromodulatory study we applied anodal transcranial direct current stimulation (tDCS) over prefrontal and parietal hubs of the frontoparietal network. tDCS selectively enhanced working memory updating, with right-hemisphere stimulation improving *Gv* but reducing *Gf* performance. Mediation analyses revealed that working memory updating mediated tDCS effects on higher cognition, yet the direction of effects varied across hemispheres and ability domains, exposing indirect, updating-driven facilitative and compensatory mechanisms within the frontoparietal network. Together, these findings bridge psychometric and neuromodulatory approaches to advance mechanistic understanding of how EFs support higher cognition.

## Introduction

Over a century of research on the structure of human intelligence has culminated in the Cattell-Horn-Carroll (CHC) model of cognitive abilities ^1^, which integrates several prominent models of cognitive abilities that have emerged in the 20^th^ century ^2–7^ and provides the dominant framework for contemporary intelligence testing. The CHC model postulates a three-stratum hierarchical organization of human cognitive abilities. At the bottom level (Stratum I), it encompasses numerous narrow abilities that reflect specific cognitive skills. These narrow abilities are grouped into broader domains at the middle level (Stratum II). These broad group factors, among others, include: fluid reasoning (*Gf*) – the capacity for abstract reasoning, solving new, complex problems using inductive and deductive reasoning; crystallized abilities (*Gc*) – breadth and depth of acquired knowledge of the language, information, and concepts of a specific culture; visual processing (*Gv*) – the ability to generate, store, retrieve, and transform complex visual stimuli; processing speed (*Gs*) – the ability to fluently perform relatively easy but attention- and focus-demanding cognitive tasks. Finally, general cognitive ability, or the *G*-factor, is positioned at the highest level of the hierarchy (Stratum III) and is thought to underlie all complex, higher-order cognitive processes ^2^.

Cognitive abilities, in general, are thought to rely heavily on executive functions (EF) as building blocks of higher cognition ^8,9^. Several research frameworks, such as the Process overlap theory (POT) ^9^ and the attention control framework ^10–12^ propose that different cognitive ability tasks tap overlapping domain-general processes that are not limited only to a particular content or specific cognitive ability, thus accounting for the positive manifold among cognitive abilities, i.e., positive correlations between various cognitive tasks that give rise to *G*-factor ^9,10^. These processes are believed to have a crucial role in overall cognitive functioning and are likely to serve as a “bottleneck” within the information processing system ^9,10^. However, since EFs are not a unitary construct, likely candidates for these domain-general processes remain debated.

The Unity-diversity model of EF ^13–15^ offers a set of viable candidates for domain-general processes subserving higher-order cognition. In its initial form, the model proposed three distinct yet related EFs – updating, inhibition, and shifting ^14^. Updating of working memory (WM) representations enables dynamic and continuous monitoring and encoding relevant information in WM and their simultaneous revision; inhibition reflects deliberate suppression and overriding of dominant, automatic, or prepotent responses, while shifting reflects the ability to efficiently shift attention from one task, operation, or mental set to another. These functions conceptually overlap with distinct yet complementary domain-general processes of maintenance and disengagement, defined within the attention control framework, which are thought to underpin performance in various cognitive tasks ^11,12^. In recent years, an alternative perspective on the Unity-diversity model has gained prominence ^16,17^. This alternative model differentiates between three orthogonal latent factors: general executive function abilities (Common EF), abilities specific to updating (Updating-specific), and shifting (Shifting-specific). Common-EF abilities are believed to reflect active creation and maintenance of task goals and their use in biasing lower-level processing, updating-specific abilities reflect selective gating of information into and out of WM, while shifting-specific abilities are reflected in flexible replacement of goals that are no longer necessary ^13,15,18^. In this model, there is no inhibition-specific factor since empirical findings showed that all variance of inhibition is captured by Common EF ability ^13,15,18^.

Previous psychometric studies have shown that EFs and cognitive abilities are closely related yet distinct constructs ^19–21^. Moreover, not all EFs have been found to correlate with cognitive abilities. For instance, some findings have shown that only updating is significantly related to both fluid and crystallized intelligence ^19^. The alternative Unity-diversity approach showed that both Common-EF and Updating-specific abilities, but not Shifting-specific abilities, were moderately correlated with full-scale IQ assessed by the Wechsler Adult Intelligence Scale (WAIS) full-scale ^18^. However, empirical evidence on the relationships between EFs and broad cognitive abilities remains largely understudied.

Although EFs are widely considered as functions of the dorsolateral prefrontal cortex (DLPFC), their unity and diversity found on a behavioral, i.e., psychometric level ^13–15^ can be observed on the neural level too. Studies have shown that solving various EF tasks consistently recruits a similar set of brain regions within the frontoparietal (i.e., central executive network, CEN) and cingulo-opercular networks ^17,22,23^. However, while the DLPFC has been widely implicated in executive control, especially working memory updating ^24^, the evidence for its role in inhibition and shifting is weaker ^25–27^. Moreover, all three EFs appear to engage both frontal as well as posterior parietal regions ^22,24,27–31^.

Converging evidence from neuroimaging studies supports a shared neural basis for EFs and cognitive abilities within the frontoparietal network. The Parieto-frontal integration theory (P-FIT), as the most parsimonious account for many of the empirical findings linking individual differences in intelligence to variations in brain structure and function ^32^, postulates that the biological basis of individual differences in human intelligence relies on large-scale neural networks which connect and integrate brain regions, including frontal, parietal, temporal, and cingulate cortices, with the critical interaction between association cortices within parietal and frontal areas which when effectively linked by white matter structures underpin individual differences in intelligence ^32^.

To better understand the functional role of EFs in higher cognition, it is essential to move beyond purely correlational approaches. While psychometric and neuroimaging methods have provided valuable insights, they are limited in their ability to directly test causal links between EFs and higher-order cognition. In this context, neuromodulation techniques, such as transcranial direct current stimulation (tDCS), offer a powerful experimental tool for examining how modulating brain activity influences both EFs and cognitive abilities. tDCS modulates cortical excitability by inducing weak electrical currents between scalp electrodes (typically 1–2 mA), which, depending on the polarity of stimulation (anodal, i.e., excitatory or cathodal, i.e., inhibitory), can depolarize or hyperpolarize neurons, thereby altering the likelihood of their firing ^33,34^. tDCS aftereffects are believed to involve NMDA receptor-mediated plasticity and resemble long-term potentiation (LTP) and depression (LTD) ^34^. Given its good safety profile ^35^ and high tolerability ^36^, tDCS has been widely used to probe brain activity in cognitive neuroscience.

Meta-analytic evidence generally supports the use of tDCS to probe EFs; however, the results are largely inconsistent across studies. Several meta-analyses have shown that tDCS over prefrontal brain areas, specifically the DLPFC, has the potential to enhance WM updating ^37,38^. Some evidence also suggests that tDCS can modulate inhibitory control, although these effects are usually task-specific and generally weaker ^39,40^ while empirical evidence on tDCS effects on shifting remains limited, with some meta-analyses showing null effects, while others show significant tDCS effects ^38,39,41^. However, posterior brain regions remain underinvestigated in tDCS research of EFs, despite converging neuroimaging evidence of their involvement in cognitive control ^22,24,27–31^.

Unlike EFs, tDCS studies on cognitive abilities remain scarce and yield inconsistent findings. For instance, Sellers et al. ^42^ found that excitatory anodal tDCS over left and right DLPFC, as well as bilateral tDCS of both areas, impaired cognitive performance on Full-scale IQ WAIS-IV, primarily due to decreased perceptual reasoning following stimulation of the right DLPFC, with a similar trend observed for left-sided and bilateral stimulation too ^42^. A subsequent study using high-definition tDCS (HD-tDCS), however, showed that stimulation of left and right DLPFC did not affect cognitive performance on Raven’s matrices ^43^.

### Current research

Here, we integrate psychometric and neuromodulation approaches to advance understanding of the role of EFs in higher-order cognition. In Study 1, we mapped the psychometric relationships between EFs – updating, inhibition, and shifting, as defined by the Unity-Diversity model ^13–15^, and four broad cognitive abilities – fluid reasoning (*Gf*), crystallized abilities (*Gc*), visual processing (*Gv*), and processing speed (*Gs*) as defined within the CHC model of intelligence ^1^. Building on the psychometric findings from Study 1 and using the same set of cognitive measures, Study 2 probed the functional role of EFs in higher cognition by targeting their shared neural substrates. Using anodal (excitatory) tDCS applied to key neural hubs of the frontoparietal network, the DLPFC, and the posterior parietal cortex (PPC), we tested if neuromodulation of EFs (updating, inhibition, shifting) would mechanistically extend to broad cognitive abilities (*Gf*, *Gc*, *Gv*, *Gs*). We tested whether the psychometric architecture linking EFs and cognitive abilities would be mirrored in tDCS-induced performance changes, thereby grounding behavioral associations in neurophysiological mechanisms.

### Study 1

The first study systematically assessed the relationship between EFs and cognitive abilities, focusing on both the model of correlated EF factors (updating, inhibition, and shifting) and an alternative EF model (Common EF, Updating-specific, and Shifting-specific), in relation to four broad cognitive ability factors (*Gf*, *Gc*, *Gv*, and *Gs*). In line with previous findings ^19^, we hypothesized that each cognitive ability would be more strongly associated with updating than with inhibition or shifting. We further expected that Common EF would correlate with all broad ability factors ^18^. However, drawing on an extensive body of research on the role of WM in *Gf* ^44–47^, we expected that Updating-specific abilities would correlate more strongly with *Gf* than other abilities. Finally, we hypothesized that Shifting-specific abilities would be unrelated to any broad factor of cognitive abilities ^18^.

## Methods

### Participants

Two hundred eighteen participants took part in Study 1 (*M* = 20.50, *SD* = 1.30, 79.8% females). The participants were undergraduate and graduate students at the University of Belgrade, and they participated in the study on a voluntary basis in exchange for course credits.

### Procedure

The test battery consisted of 12 short cognitive ability tests (three per each cognitive ability factor *Gf*, *Gc*, *Gv*, and *Gs,* half verbal and half nonverbal/spatial) and 6 executive function tasks (two per each executive function – updating, inhibition, shifting, half verbal and half nonverbal/spatial). The tasks measuring executive functions were programmed and administered in OpenSesame. Tests of *Gf*, *Gc*, and *Gv* were computer-administered while participants completed *Gs* measures in paper-and-pencil form.

Cognitive tests were administered in two counterbalanced testing blocks: one for the assessment of executive functions and another for the assessment of cognitive abilities. For practical reasons, the subjects completed *Gs* tests in a paper-and-pencil format at the beginning of the test session, after which the remaining cognitive tests were computer-administered. Before each test, the participants had a short practice session to familiarize themselves with the tasks they were about to solve. The assessment session lasted approximately 60 minutes.

### Cognitive tasks

#### Updating

The updating was assessed using verbal and spatial 3-back tasks, each consisting of one practice and five testing blocks ^48^. In verbal 3-back tasks (Figure 1A), ten consonants of the Latin alphabet (S, F, H, C, G, R, N, B, V, T) printed in black ink and presented on a white background were used as stimuli. In each block, 32 letters (25% targets) were presented successively in fixed, pre-randomized order. Each letter was presented with a fixed interval of 1000ms (ISI = 500ms), and participants were instructed to respond via keypress when the presented letter was identical to the one shown three steps before. In the spatial 3-back task (Figure 1B), participants were presented with a 3×3 matrix on a white background with a black square quasi-randomly appearing in one of nine matrix fields for 1000ms (ISI = 500ms). Each block had 32 stimuli (25% targets), and participants were instructed to respond only when the black square appeared at the identical position within the matrix as three steps before.

**Figure 1.**
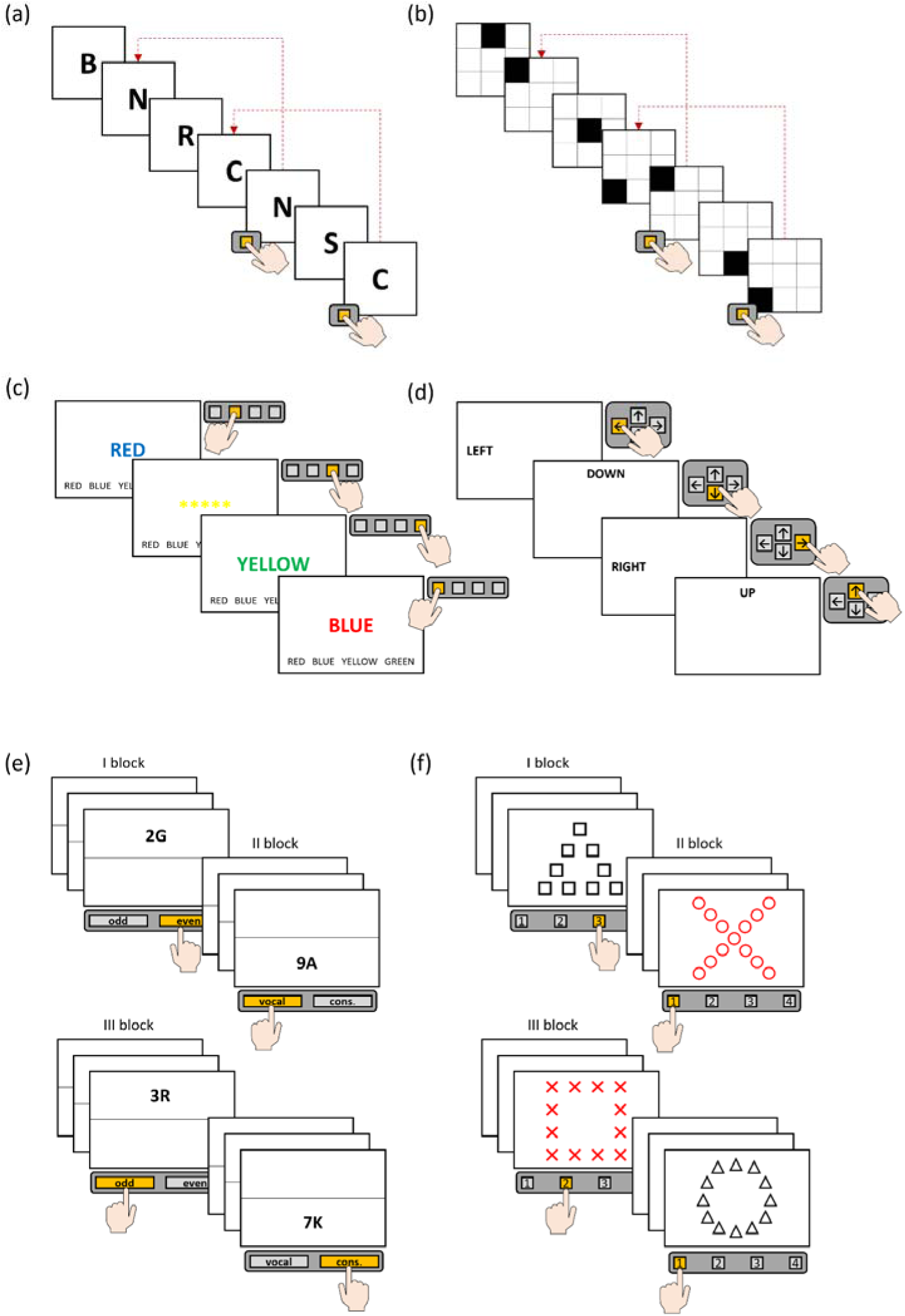
Tasks of executive functions. – (A) Verbal 3-back; (B) Spatial 3-back; (C) Verbal Stroop task; (D) Spatial Stroop task; (E) Number-letter; (F) Local-global

#### Inhibition

The inhibition was assessed via computerized verbal and spatial Stroop tasks ^49^. In the verbal inhibition task (Figure 1C), participants were sequentially presented with a list of congruent (e.g., word “red” printed in *red*), neutral (e.g., asterisks printed in red), and incongruent stimuli (e.g., word “red” printed in *yellow*). Participants were instructed to indicate the color in which each stimulus was presented via a key press. The measure of inhibition was calculated by subtracting the average reaction times (RTs) in incongruent trials from the average RTs in neutral trials. In the spatial Stroop task (Figure 1D), participants were sequentially presented with words indicating spatial positions (*up*, *down*, *left*, *right*) in congruent and incongruent positions on the screen. They were instructed to inhibit spatial interference (the position of the words on the screen) and only respond to the verbally presented information. The score was calculated by subtracting average RTs for incongruent opposite stimuli (e.g., the word *down* appearing at the *top* of the screen) from the average RTs in congruent conditions. In both tasks, the order of stimuli within each form was pre-randomized and thus fixed for all participants.

#### Shifting

The shifting was assessed by the verbal Number-letter and nonverbal Local-global task ^14,50^. Both tasks had a similar structure consisting of two mono-blocks and one hetero-block. In the Number-letter task (Figure 1E), participants were sequentially presented with number-letter pairs. During the first block, number-letter pairs were shown in the upper half of the screen, and the participants were instructed to indicate if the number was even or odd. During the second block, the number-letter pairs were presented in the lower half of the screen, and the participants were instructed to indicate if the letter was a vowel or a consonant. Finally, in the third block, number-letter pairs were presented either in the screen’s upper or lower half, requiring odd/even or vowel/consonant responses, respectively. In the Local-global task (Figure 1F), the participants were sequentially presented with Navon’s figures ^51^, in which the contours of larger global figures were outlined by smaller local figures. In the first block, the figures were presented in black, and participants needed to indicate the number of lines that a global figure consists of. In the second block, the figures were shown in red, and participants were instructed to indicate the number of lines each of the local figures consists of. Finally, in the third block, figures were presented in either black or red, and participants needed to respond to either global or local stimulus characteristics depending on the figure’s color. The cost of shifting, i.e., the slowdown in the RTs in the third block compared to the first two blocks, was calculated by subtracting the average RTs in the third block from the average RTs in the first two blocks.

#### Fluid abilities (*Gf*)

To assess *Gf*, the Matrix reasoning test (MTRX) ^52–54^, Fluid Analogies (FAL) ^52^, and the short form of Raven’s Progressive Matrices (RPM) were used ^55^. In the MTRX test (Figure 2A), participants were presented with 16 matrices or series of figures with one missing element, and instructed to select the element that best complements the matrix among the six available options. The time limit in this test was five minutes. In the FAL test (Figure 2B), participants were presented with 31 analogies and instructed to choose among five response options and select the one in which the relationship between two concepts was the same as in the target analogy. Participants had five minutes to complete the test. The RPM (Figure 2C) consisted of 18 matrices with figural elements in which the bottom right corner is missing. The participant’s task was to figure out the pattern and select the image that best complements the matrix from among five response options. Participants had six minutes to complete the test.

**Figure 2.**
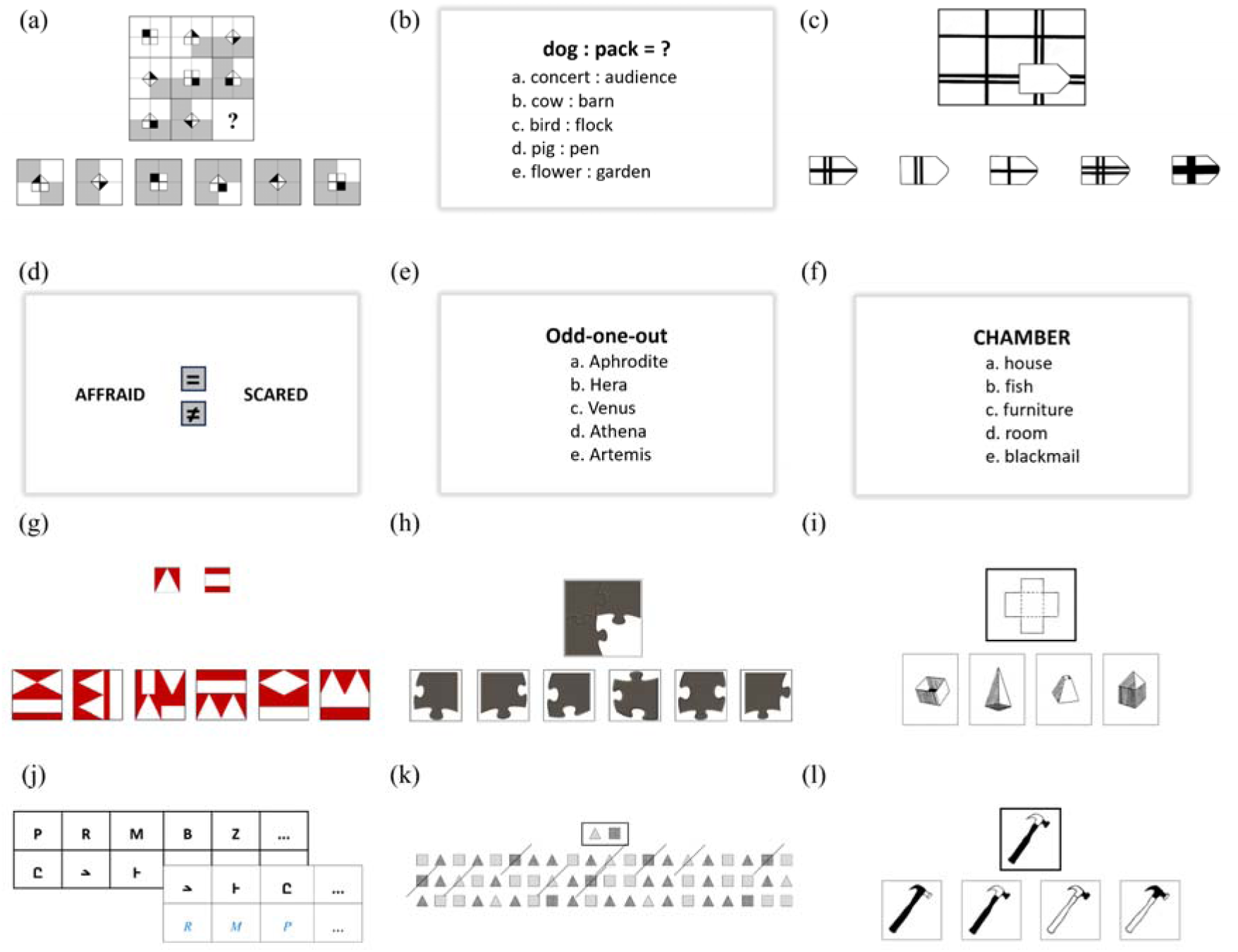
Cognitive ability tests. – (a) Matrix test (MTRX); (b) Fluid analogies (FAL); (c) Raven’s progressive matrices - short form (RPM); (d) Synonym-antonym (S-A); (e) Crystallized associations; (f) Lexical knowledge test (GSN); (g) Mosaic (MCS); (h) Puzzles (PZL); (i) Spatial ability test (IT2); (J) Symbol (SYM); (K) Visual search (VS); (L) Identical figures (IT1)

#### Crystallized abilities (*Gc*)

To assess *Gc,* three verbal tests were used – the Synonyms-Antonyms (S-A) ^52^, Crystallized associations test (CA) ^52^, and Lexical knowledge test (GSN) ^56^. In the Synonyms-Antonyms test (Figure 2D), participants were sequentially presented with 54 word pairs. Their task was to indicate as fast as possible if the presented words have the same or opposite meaning. Participants had 90 seconds to complete the test. In the Crystallized associations test (Figure 2E), participants were presented with 32 items, each consisting of five response options – four of which belong to the same category and one that does not fit with the others. The participants were instructed to select the option that does not belong to a given group. CA items were based on the following domains of knowledge: history of science and discovery, politics, sports, science, medicine/biology, history, geography, literature, visual arts, music, film, games, fashion, finances, and pop culture. The time limit for this test was four minutes. GSN test (Figure 2F) consisted of 39 items in which participants were presented with a target word and five response options and instructed to select the word with the most similar or the same meaning as the target word. The time limit in this test was two minutes.

#### Visual processing (*Gv*)

To assess *Gv*, three figural, non-verbal tests were used – Mosaic (MSC) ^52^, Puzzles (PZL) ^52^, and the spatial ability IT2 test ^56^. In the MSC, participants were presented with the figural element/s (1-3 squares with a specific pattern) and six mosaics. They were instructed to select the mosaic that could be completed by combining given figural elements (Figure 2G). The test consisted of 35 items, and participants had seven minutes to complete it. In the Puzzle test, participants were presented with 36 puzzles of varying complexity, each lacking one element that would correctly complete it (Figure 2H). Their task was to find the element that adequately completes the puzzle among the six response options. The time limit in the PZL test was seven minutes. IT2 consisted of 39 items in which participants were presented with a target geometrical figure and four unfolded figure cutouts. Participants were instructed to select the response option, which would result in a target figure when folded in specified places (Figure 2I). The time limit for this test was 10 minutes.

#### Processing speed (*Gs*)

To assess *Gs*, we used the verbal Symbol test (SYM) ^52^, the nonverbal Visual search (VS) ^52^, and the Identical figures (IT1) test ^56^. In the SYM test (Figure 2J), participants were given a “codebook” containing ten unfamiliar symbols associated with the ten letters of the Latin alphabet. Their task was to use the “codebook” and, in sequence, transcribe as many letters as possible below each corresponding symbol in 60 seconds. In the VS test (Figure 2K), the participants were presented with a long list of figures that vary in shape (squares and triangles) and color (grey and black). Their task was to go through the list in sequence and simultaneously find and cross off as many target stimuli as possible in 60 seconds. IT1 test (Figure 2L) consisted of 39 items in which participants were presented with an image of a target tool and four response options. They were instructed to select the image of a tool identical to the target image. The time limit for this test was four minutes.

## Data analysis

Data were analyzed using IBM SPSS Statistics and Mplus 7. To examine the structural validity of EFs and cognitive abilities, we conducted a series of Confirmatory Factor Analyses (CFA) with robust MLMV estimation. For EFs, we first tested the correlated three-factor model in which each task loaded only on its respective factor, with correlations between factors freely estimated. Next, we tested an alternative bifactor model, with all tasks loading on the Common EF factor, and updating and shifting tasks additionally loading on the Updating- and Shifting-specific factors, respectively. To isolate the unique variance in the specific factors, correlations among the three factors were fixed to zero. For cognitive abilities, we tested both a group-factor model and a hierarchical model. In the group-factor model, each test loaded solely on its respective factor, with correlations among factors freely estimated. In the hierarchical model, a higher-order *G* factor was specified to account for the covariation among the group factors. Finally, the latent relationships between EFs and group factors of cognitive abilities were tested separately for the model of correlated EFs and the alternative/bifactor EF model using structural equation modeling (SEM) by freely estimating covariation between each pair of latent factors. The following indices of model fit and cut-off criteria were used: χ^2^ test, Comparative fit index – *CFI* ≥ 0.95, Tucker-Lewis fit index *– TLI* ≥ 0.95, Root Mean Square Error of Approximation *– RMSEA* ≤ 0.06, and Standardized Root Mean Square Residual *– SRMR* ≤ 0.06 ^57^. For models with suboptimal fit, modification indices were inspected to identify sources of misfit, and models were respecified to achieve satisfactory fit.

## Results

Descriptive statistics and correlations for EF tasks are shown in Table S1, and for cognitive ability tests in Table S2, while correlations between EF tasks and tests of cognitive abilities are shown in Table S3 of the Supplementary material.

For EF, both the model of three distinct yet correlated factors [χ^2^ = 6.783, *p* = .341, *CFI* = .995, *TLI* = .987, *RMSEA* = .024 (90% CI: .000-.094), *SRMR* = .029] and bifactor model^1^ [χ^2^ = 7.045, *p* = .424, *CFI* = 1.000, *TLI* = .999, *RMSEA* = .005 (90% CI: .000-.084), *SRMR* = .029], showed excellent fit (Figure 2). The pattern of latent correlations was largely comparable to previous findings, with inhibition being highly correlated to both updating and shifting, with a somewhat lower correlation between the latter two (Figure 3A). The bifactor model of EFs corroborated the alternative view on three EFs, with a Common EF factor fully accounting for the variance of inhibition with separate Updating- and Shifting-specific latent factors (Figure 3B).

**Figure 3.**
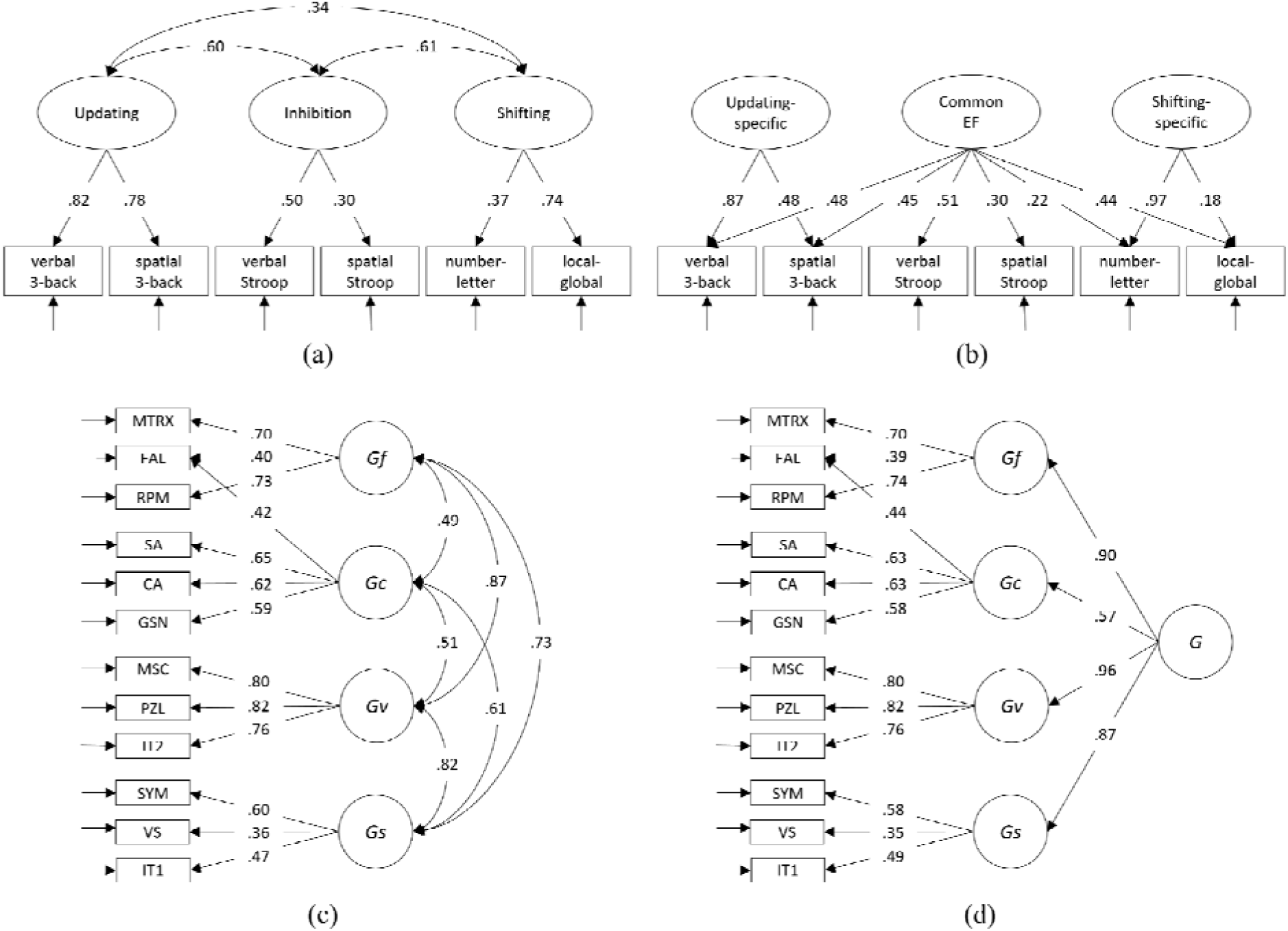
(A) Model of correlated executive functions; (B) Bifactor/nested model of EF; (C) Group-factor model of cognitive abilities; and (D) Hierarchical model of cognitive abilities

For cognitive abilities, both the group factor model [χ^2^ = 107.147, *p* < .001, *CFI* = .907, *TLI* = .872, *RMSEA* = .075 (90% CI: .056-.094), *SRMR* = .063] and hierarchical model [χ^2^ = 112.085, *p* < .001, *CFI* = .903, *TLI* = .871, *RMSEA* = .075 (90% CI: .057-.094), *SRMR* = .064], showed marginally acceptable fit. To improve model fit, FAL was allowed to load on both *Gf* and *Gc* due to its verbal content. After this modification, both the group factor model [χ^2^ = 89.609, *p* < .001, *CFI* = .933, *TLI* = .906, *RMSEA* = .064 (90% CI: .044-.085), *SRMR* = .056] and hierarchical model [χ^2^ = 92.302, *p* < .001, *CFI* = .932, *TLI* = .908, *RMSEA* = .064 (90% CI: .043-.083), *SRMR* = .056], showed satisfactory fit. All cognitive ability tests significantly loaded on their respective factors (Figure 3C). As expected, two figural reasoning tests proved to be fairly good indicators of *Gf*, while the analogical reasoning had approximately equal loadings on both *Gf* and *Gc*. Each of the three cognitive tests tapping verbal knowledge loaded highly on the *Gc*, while three tests measuring visuospatial processing showed substantial and relatively uniform *Gv* loadings. Finally, three tests measuring processing speed significantly loaded on *Gs*. Among individual cognitive ability tests, the highest *G*-loadings were observed for *Gv* measures [range: .741 (IT2) – .791 (PZL)], followed by *Gf* tests [range: .635 (FAL) – .665 (RPM)], while lower *G*-loadings were observed for *Gs* measures [range: .289 (VS) – .494 (SYM)] and *Gc* [range: .259 (GSN) – .493 (CA)]. Expectedly, correlations between the four latent factors were all positive and moderate-to-high in magnitude (Figure 3C). All latent dimensions showed substantial general factor loadings (Figure 3D), with somewhat lower general factor loading for *Gc* compared to other ability factors, likely due to the fact that nonverbal tests were overrepresented in the test battery.

To examine relationships between the two domains, we tested unified models linking latent cognitive ability factors (group-factor model) to EF factors – updating, inhibition, and shifting [χ^2^_(113)_ = 182.469, *p* < .001, *CFI* = .904, *TLI* = .870, *RMSEA* = .053 (90% CI: .038-.067), *SRMR* = .058], as well as to Common EF, Updating-specific, and Shifting-specific latent factors [χ^2^_(112)_ = 178.655, *p* < .001, *CFI* = .908, *TLI* = .874, *RMSEA* = .052 (90% CI: .037-.066), *SRMR* = .055]. Latent correlations (Table 1) revealed that *Gf* was most strongly linked to updating, showed also association with inhibition, and was unrelated to shifting. Both *Gv* and *Gs* were significantly correlated with all three EF domains, with the strongest associations observed for inhibitory control. By contrast, *Gc* showed no significant relationship with any EF domain.

**Table 1.** Psychometric relationship between EFs and cognitive abilities on the latent level.

|  | *Three-factor model of EF* |  |  | *Bifactor/nested model of EF* |  |  |
| --- | --- | --- | --- | --- | --- | --- |
|  | *Updating* | *Inhibition* | *Shifting* | *Common EF* | *Updating specific* | *Shifting specific* |
| *Gf* | .447*** | .379** | .074 | .387** | .249* | -.159 |
| *Gc* | .097 | .257<sup>†</sup> | .056 | .303* | -.085 | -.192 |
| *Gv* | .345*** | .518** | .261** | .491*** | .112 | -.001 |
| *Gs* | .428*** | .591** | .496*** | .645*** | .132 | .018 |

Note. <sup>†</sup> $p < .10$ ; \* $p < .05$ ; \*\* $p < .01$ ; \*\*\* $p < .001$

The Common EF latent factor had moderate to strong associations with the latent factors of all four cognitive abilities, fully accounting for the relationship between inhibition and cognitive abilities. Notably, *Gf* was moderately correlated with the Common EF latent factor and uniquely emerged as the only cognitive ability factor significantly associated with the Updating-specific factor. Shifting-specific abilities, on the other hand, were not related to any cognitive ability factor.

## Discussion

The present findings extend previous evidence on the psychometric architecture of the associations between prominent conceptualizations of the EFs ^13,15,58^ and cognitive abilities ^1^. Our results support the view that EFs and cognitive abilities are closely related yet clearly separable constructs ^19–21^, with heterogeneous associations between them. Consistent with prior studies ^19^, *Gf* showed the strongest link with updating, adding to the body of evidence showing that the dynamic monitoring and revision of relevant information in WM is one of the pillars of reasoning abilities. Additionally, *Gf* was also moderately related to inhibition, suggesting an important role of interference control in effective reasoning. In contrast, *Gv* demonstrated significant associations with all three EF domains, most strongly with inhibition, moderately with updating, and only weakly related to shifting abilities, suggesting that visuospatial manipulation critically relies on effective interference control as well as monitoring and revising goal-relevant information in WM. Similarly, all three executive functions correlated with *Gs*, which emerged as a group factor attaining, on average, the highest correlations with EFs, being most closely linked to inhibition and shifting, and achieving somewhat lower correlation with updating. This finding suggests that processing speed is strongly supported by the ability to rapidly suppress irrelevant information and flexibly shift attention, highlighting its close ties to fast and efficient executive control, with less reliance on updating processes. Contrary to previous findings ^19,20^, we found no significant association between *Gc* and any domain of EFs in particular, indicating that *Gc*, as a store of acquired knowledge, is less reliant on domain-general executive processes and more dependent on domain-specific processes and long-term memory systems.

The alternative bifactor model of EFs offered a more nuanced understanding of the distinctive and shared cognitive mechanisms underpinning cognitive abilities, thereby building upon and extending previous findings showing that both Common EF and Updating-specific abilities significantly correlate with IQ ^18^. However, our results showed that different cognitive abilities rely on EFs to varying extents. Specifically, *Gv* and *Gs* showed strong associations with the Common EF factor and were unrelated to Updating- or Shifting-specific processes highlighting the importance of attention maintenance, which is considered the core of the Common EF factor, in both visuospatial abilities and fluent and efficient information processing. These findings suggest that effective goal maintenance, sustained attention, and interference control are essential for maintaining and manipulating visual information, as well as for the rapid activation of task rules, efficient visual scanning, and resistance to distractors, which are reflected in *Gv* and *Gs,* respectively. Moreover, *Gv* and *Gs*, judging by the magnitude of their correlations with the Common EF factor, appear to be more reliant on general EF abilities than *Gf* and especially *Gc*, further confirming that, in contrast to other ability factors, crystallized abilities rely more on domain-specific than domain-general executive processes. *Gf*, however, emerged as the only ability factor correlated with both general attention control processes as well as executive processes directly supporting updating of representations in WM. This finding is consistent with decades-old proposals of the significance of WM capacity in fluid abilities as well as in *G* ^44–47^. This dual association with both general attention control and abilities underlying selective gating of information in and out of WM underscores the complex cognitive architecture underlying reasoning ability and appears to be a key feature distinguishing *Gf* from other broad cognitive abilities.

Lastly, and particularly relevant for Study 2, all instruments demonstrated good psychometric properties, including structural validity (model fits) and proved to be sufficiently challenging, yielding adequate variability in performance, thereby ensuring their suitability for use in subsequent tDCS experiments.

### Study 2

In Study 2, we examined the short-term effects of anodal tDCS over the frontoparietal functional network on EFs (updating, inhibition, and shifting), and broad cognitive abilities (*Gf*, *Gc*, *Gv*, and *Gs*). The core hypothesis was that anodal tDCS would selectively increase activity in the targeted area(s), thus highlighting their functional contribution to the investigated cognitive processes. We targeted two central hubs of the frontoparietal network – DLPFC and PPC. To account for hemispheric specialization and the task domain of assessment, both left and right hemispheres were stimulated. Cognitive performance was assessed with validated cognitive tests described in Study 1. Given the well-established role of these brain regions in EFs ^17^ and cognitive abilities ^32^, tDCS applied to DLPFC should broadly enhance cognitive performance, and PPC stimulation should have similar effects given its considerable involvement in both EFs and cognitive abilities. Alongside standard single-locus anodal tDCS, we introduced a novel exploratory condition: fronto-parietal (FP) stimulation of both DLPFC and PPC. Since anodal FP tDCS was not previously used in the context of cognition, we expected its effects to approximate those of single-locus protocols while potentially offering broader network-level effects.

Building on Study 1, we hypothesized tDCS to exert concurrent effects at corresponding brain sites, such that tDCS-induced changes in EFs would drive aligned changes in the associated cognitive abilities (*Gf*, *Gv*, and *Gs*) as a function of their psychometric relationships. By contrast, we anticipated little to no effect on *Gc*, reflecting its low reliance on domain-general EFs. Crucially, we hypothesized that tDCS effects on cognitive abilities at a given locus would be mediated by stimulation-induced changes in EF at that same locus.

## Method

### Participants

Forty-eight neurotypical adults (age range: 21-35, *M* = 26.10, *SD* = 4.77 years, 24 females) participated in the experiment. Participants were assigned to one of two groups: left hemisphere tDCS (*n* = 24) or right hemisphere tDCS (*n* = 24). An a priori power analysis conducted in G*Power indicated that a minimum of 23 participants was required to detect a moderate effect (*d* = .50) with 80% power at an alpha level of .05, assuming a correlation of *r* = .50 between repeated measures.

All participants met standard tES inclusion/exclusion criteria ^35^, and reported no history of psychiatric or neurological conditions, acute or chronic skin conditions, or the use of psychoactive substances or medication. All were native speakers with normal or corrected-to-normal vision. Written informed consents were obtained prior to participation. The study adhered to the Declaration of Helsinki and was approved by the Institutional Ethics Board.

### Experimental design

The study employed a fully-repeated, cross-over, sham-controlled design with two groups (left *vs.* right hemisphere). Each participant underwent four stimulation conditions in a counterbalanced order: 1) DLPFC, 2) PPC, 3) FP (simultaneous DLPFC-PPC stimulation), and 4) sham, with half of them receiving tDCS over the left (L-DLPFC, L-PPC, L-FP) and the other half over the right hemisphere (R-DLPFC, R-PPC, R-FP). The effects of tDCS were assessed immediately after stimulation (so-called *offline* protocol) using parallel forms of tests measuring four cognitive abilities (*Gf*, *Gc*, *Gv*, *Gs*) and three EFs (updating, inhibition, shifting).

### Transcranial direct current stimulation (tDCS)

All stimulation procedures and protocols followed safety guidelines for the use of tDCS in human studies ^35^. The stimulation was delivered via an isolated constant current stimulator (STMISOLA, BIOPAC Systems, Inc., USA) controlled by “CED 1401 Plus” using the “CED Signal” software (Cambridge Electronic Design, Cambridge, UK). Three 5×5 cm rubber electrodes were inserted into saline-soaked sponge pockets and positioned on the participant’s head in each session: two over cortical target sites and one over the cheek. In the left-hemisphere group, electrodes were placed over F3 (L-DLPFC) and P3 (L-PPC) according to the 10–20 EEG system, while in the right-hemisphere group, they were positioned over F4 (R-DLPFC) and P4 (R-PPC). In both groups, the return electrode (i.e., cathode) was placed on the contralateral cheek. In the single-locus conditions, only one target electrode (F3 or P3; F4 or P4) served as the active anode, while in the FP condition, both DLPFC and PPC electrodes acted as active anodes simultaneously. In the single-locus conditions, stimulation was delivered at 1.8 mA, corresponding to a current density of 0.072 mA/cm². In the FP condition, current intensity was set to 0.9 mA at each anode (F3/P3 or F4/P4), yielding a total of 1.8 mA at the return electrode. This resulted in a current density of 0.036 mA/cm² for each anode and 0.072 mA/cm² for the cathode. In each stimulation condition, current was applied for 19 minutes with 30-second ramp-up and ramp-down periods at the beginning and end of stimulation. In the sham condition, current was delivered only during the first and last 60 seconds (30-second ramp up/down), to mimic the cutaneous sensations of real tDCS.

### Outcome measures

Participants completed 14 short cognitive tests identical to those used in Study 1, half verbal and half nonverbal (spatial). Specifically, two tests were administered for each of the four broad cognitive ability factors: *Gf* (MTRX, FAL), *Gc* (S-A, CA), *Gv* (MSC, PZL), *Gs* (SYM, VS); as well as two tasks for each of the three EFs: updating (Verbal and Spatial 3-back), inhibition (Verbal and Spatial Stroop task), and shifting (Letter-number and Local-global). All cognitive ability tests were administered with the same time limit as in Study 1, with the exception of the S-A test, which was shortened to 60 seconds to increase its difficulty and prevent potential ceiling effects. For cognitive ability tests, parallel forms were developed, while for EF measures, each parallel form employed a different but fixed quasi-randomized order of stimuli.

### Procedure

Each participant completed four sessions, with a counterbalanced order of stimulation conditions: DLPFC, PPC, FP, and sham. Sessions were scheduled at least two weeks apart, and participants were tested at approximately the same time of day. Following the 20-minute tDCS protocol, participants completed the cognitive assessment. Tests were organized into two experimental blocks: cognitive ability tests and EF tasks. The order of blocks was counterbalanced across participants and stimulation conditions, while the sequence of tasks within each block was pre-randomized. Parallel forms of each task were also counterbalanced across participants and conditions. The participants were blind to stimulation conditions, and a two-experimenter procedure was implemented to maintain integrity of blinding. Each session lasted about 2h in total, with post-stimulation cognitive testing of approximately 60 minutes.

### Data analysis

Data analyses were conducted in IBM SPSS v21 and R v4.5.1. The equivalence between the four test forms of each cognitive task was assessed by the intraclass correlation coefficients (*ICC*; Two-way random effect method, absolute agreement type) for each outcome measure separately. To evaluate reliability of measurement, Cronbach’s alpha indices of internal consistency were computed for the *Gf*, *Gc*, and *Gv* tests. For the remaining measures, namely *Gs* and EFs, Cronbach’s alpha was not applicable because these tests/tasks are not composed of multiple items in the conventional sense. Instead, reliability was conservatively estimated by calculating the proportion of variance in each test form (*R*^2^) that was accounted for by the remaining test forms.

To provide a conventional test of stimulation effects, a series of linear mixed models was performed with CONDITION (DLPFC/PPC/FP *vs.* sham) and type of TEST as fixed factors, and participant ID as a random factor (e.g., *updating_score ∼ condition * test + (1 | ID)*). Planned contrasts were run separately for each stimulation site (*DLPFC vs. sham, PPC vs. sham, FP vs. sham*) and for each cognitive construct (updating, inhibition, shifting; *Gf*, *Gc*, *Gv*, *Gs*). Models were fitted using *lme4* (v1.1-37) and *p*-values for fixed effects obtained with *lmerTest* (v3.1-3). Effect sizes are reported as partial eta-squared (η_p_ ^2^) together with exact *p*-values.

To obtain a single score for each construct per condition, test scores were first examined for CONDITION × TEST interactions, then z-standardized, and finally aggregated within each construct (*Gf, Gc, Gv, Gs, updating, inhibition, shifting*). To isolate within-subject changes in performance, we computed deviation (gain) scores for each active condition via person-mean centering, i.e., by subtracting each participant’s within-person mean (average of active and sham) from their active score (e.g., *PPC_Gf_agregated_score_ – Mean(PPC_Gf_agregated_score_ sham_Gf_agregated_score_*). These construct-level gain scores were z-standardized and used in the mediation models.

Mediation analyses were performed using the *mediation* R package to test whether stimulation effects on cognitive abilities were mediated by tDCS-induced changes in EFs. For each tDCS contrast (DLPFC *vs.* sham, PPC *vs.* sham, FP *vs.* sham) × EF mediator (*updating, inhibition,* and *shifting* gains) × ability (*Gf, Gc, Gv*, and *Gs* gains), we fitted standard mediation models: the *a-path* (effect of tDCS on EF mediator), the *c-path* (total effect of tDCS on ability), the *b-path* (effect of EF mediator on ability controlling for tDCS), and the *c*′*-path* (direct effect of tDCS on ability controlling for EF mediator). Indirect effects (a×b) of tDCS on ability via EF mediator were also estimated. Statistical inference for indirect effects was based on a quasi-Bayesian Monte Carlo procedure with 5000 simulations.

## Results

Cognitive ability tests demonstrated good internal consistency, with Cronbach’s alphas ranging from .58 to .93 across tests and forms, with a median value of .86 (Table S4). Consistency across forms for cognitive ability measures, as indexed by *ICCs*, was particularly high, with most values exceeding .90. Relatively high consistencies between forms were also found for EFs, despite generally lower reliability indices, ranging from .21 to .73, with a median value of .54. Especially low reliability indices were found for RT-based measures of inhibition and shifting. However, overall high *ICC* values indicated high cross-form consistency and supported the interchangeability of forms for each construct measured. Descriptive statistics for all outcome measures by stimulation conditions are presented in Table S5. All cognitive measures demonstrated adequate difficulty and variability across stimulation conditions.

Conventional mixed-effects tests of the main effects of CONDITION, TEST, and CONDITION × TEST interactions are reported in the Supplement (Table S6). Since, no CONDITION × TEST interaction was observed throughout, we analyzed tDCS effects using construct-level deviation scores (i.e., gains). Accordingly, the results below focus on the effects of tDCS on EF (*a-paths*) and cognitive abilities (*c-paths*) within a unified mediation framework (Figure 4), with direct (*c*′*-path*) and indirect (*a×b*) effects reported in subsequent mediation results.

**Figure 4.**
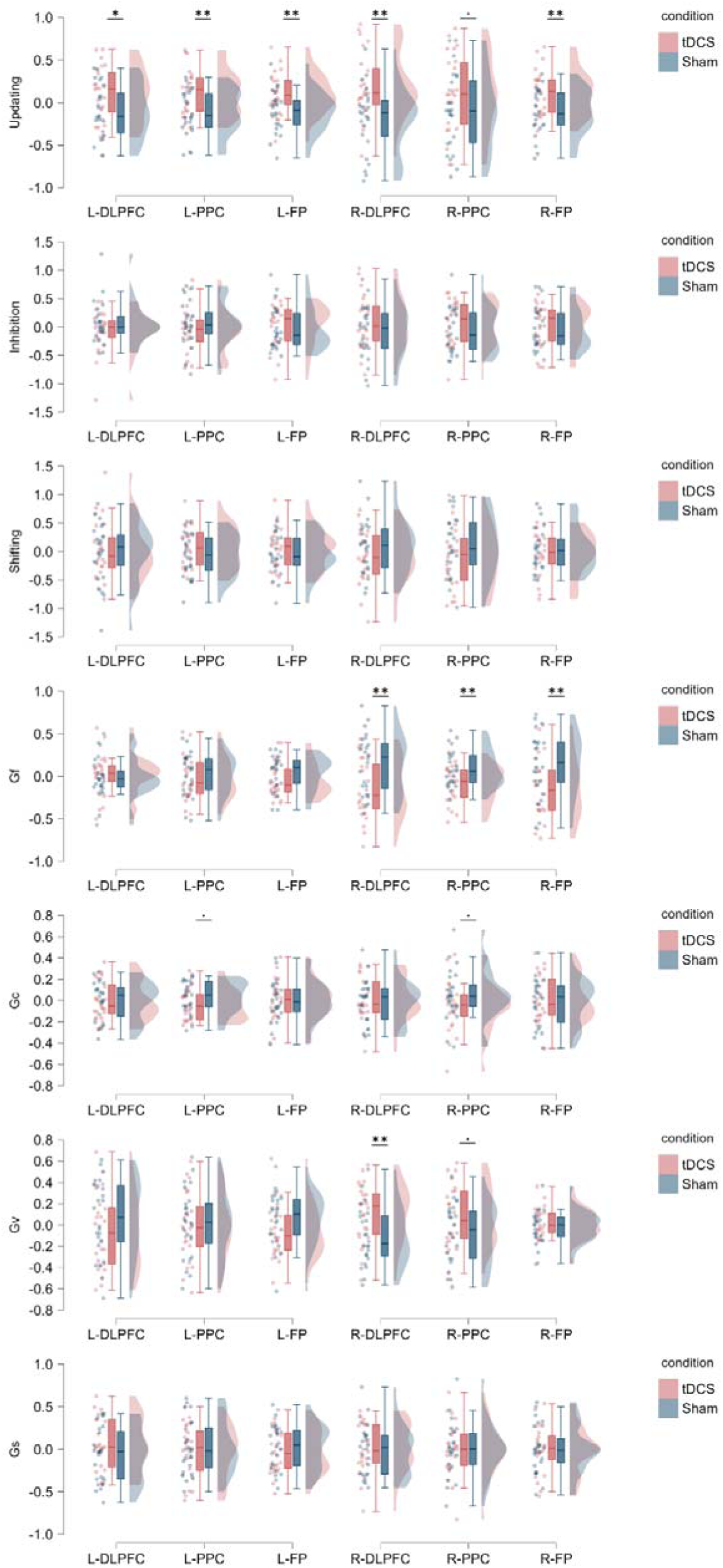
**The effects of tDCS on EFs and cognitive abilities; **p <.01, *p <.05, . p <.10**

For EFs, updating performance was enhanced by stimulation of L-DLPFC (*B* = 0.643, *SE* = 0.270, *p* = .021), L-PPC (*B* = 0.784, *SE* = 0.262, *p* = .004), as well as L-FP stimulation (*B* = 0.784, *SE* = 0.262, *p* = .004). In the right hemisphere, facilitatory effects on updating were found for R-DLPFC (*B* = 0.888, *SE* = 0.255, *p* = .001) and R-FP stimulation (*B* = 0.790, *SE* = 0.262, *p* = .004), with a similar trend for R-PPC stimulation (*B* = 0.461, *SE* = 0.278, *p* = .104). No reliable effects were found for inhibition or shifting, under either left (all *p*-values > .149) or right hemisphere conditions (all *p*-values > .171).

For cognitive abilities, R-DLPFC stimulation improved *Gv* performance (*B* = 0.735, *SE* = 0.265, *p* = .008), with a marginally significant effect in the same direction for R-PPC (*B* = 0.551, *SE* = 0.274, *p* = .050). By contrast, *Gf* performance was significantly disrupted following R-DLPFC (*B* = -0.811, *SE* = 0.261, *p* = .003), R-PPC (*B* = -0.792, *SE* = 0.262, *p* = .004), as well as R-FP stimulation (*B* = -0.750, *SE* = 0.264, *p* = .007). Trend-level reductions in *Gc* performance were observed for both R-PPC (*B* = -0.471, *SE* = 0.277, *p* = .096) and L-PPC (*B* = -0.470, *SE* = 0.278, *p* = .097). No effects of stimulation were found for *Gs*.

Mediation analyses revealed that updating served as the principal pathway through which tDCS altered cognitive abilities performance (Figure 5). Specifically, after controlling for updating gains, negative direct effects on *Gv* emerged for L-DLPFC (*B* = -0.706, *SE* = 0.252, *p* = .007) and L-FP (*B* = -0.730, *SE* = 0.282, *p* = .013), with a similar trend for L-PPC (*B* = -0.508, *SE* = 0.274, *p* = .070). In parallel, all three sites showed significant positive indirect effects via updating (L-DLPFC: *B* = 0.352, 95%CI [0.056, 0.726], *p* = .014; L-PPC: *B* = 0.399, 95%CI [0.108, 0.774], *p* = .002; L-BL: *B* = 0.310, 95% CI [0.058, 0.652], *p* = .007), revealing a suppressor mediation pattern in which updating-related improvements masked adverse direct effects on *Gv.* Conversely, in the right hemisphere, partialling out updating eliminated *Gv* enhancement following R-DLPFC (*B* = 0.278, *SE* = 0.258, *p* = .287) and R-PPC stimulation (*B* = 0.324, *SE* = 0.245, *p* = .191). Here, positive indirect effect via updating was significant for R-DLPFC (*B* = 0.456, 95% CI [0.157, 0.834], *p* < .001) with a similar trend observed for R-PPC as well (*B* = 0.229, 95% CI [-0.039, 0.562], *p* = .095).

**Figure 5.**
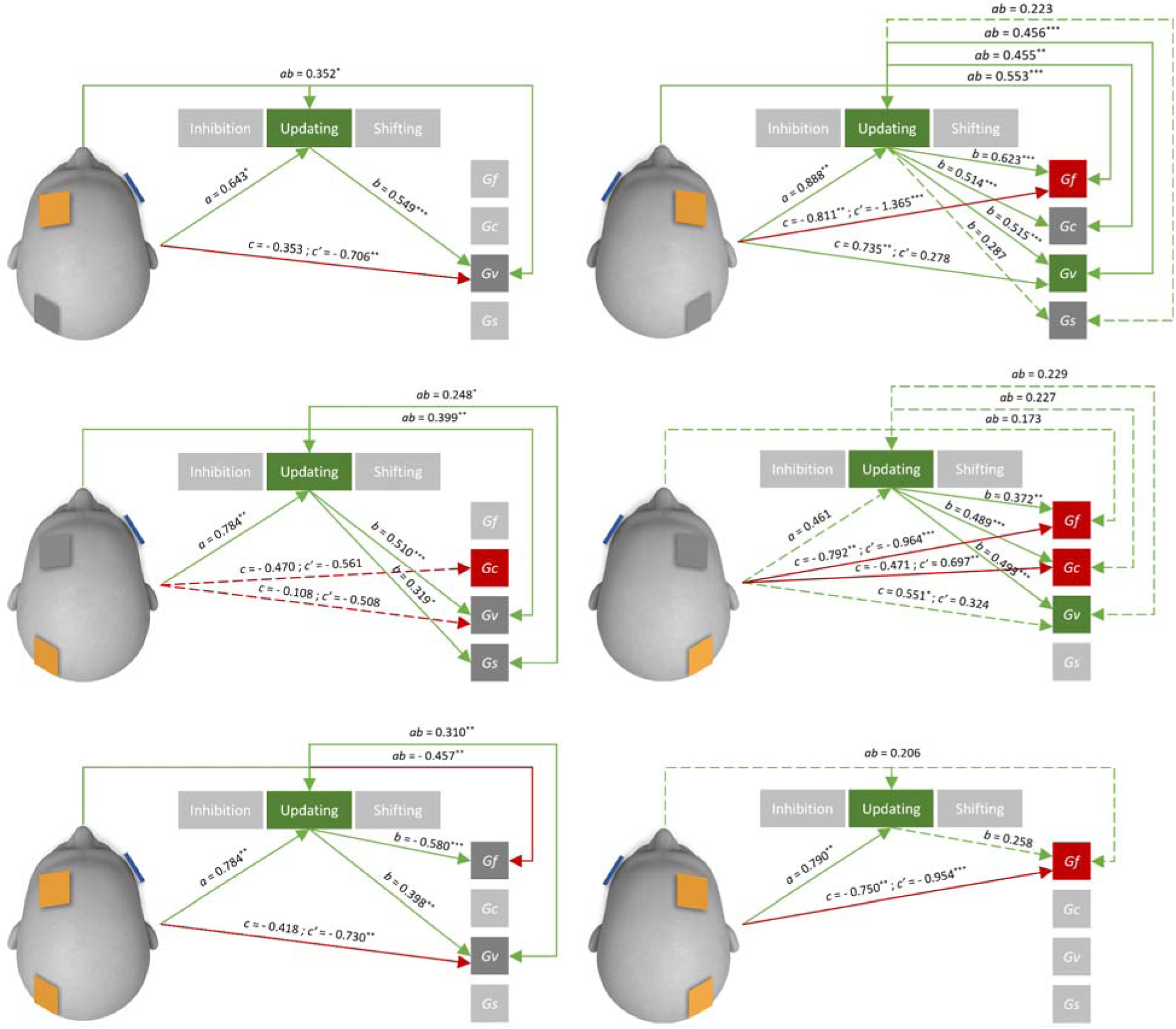
Mediation models of tDCS effects on executive functions (EFs) and cognitive abilities, with Updating as mediator. tDCS conditions are illustrated schematically, showing the positively charged (orange), inactive (grey), and negatively charged return (blue) electrodes. Solid arrows denote significant paths (**\*\*\****p* <.01, **\*\****p* <.01, **\****p* <.05), whereas dashed arrows indicate trend-level effects (*p* <.10). *B* coefficients are displayed above each corresponding path. Green and red arrows represent positive and negative effects, respectively, across the following model components: the effect of tDCS on EFs (*a*-path), the effect of EFs on cognitive abilities (*b*-path), the direct effect of tDCS on cognitive abilities (*c*′-path), and the total effect (*c*-path). EF and cognitive abilities are color-coded to indicate net positive (green), net negative (red), or null (grey) effects. The indirect (*ab*) effects highlight that the influence of tDCS on cognitive abilities was mediated by Updating.

For *Gf,* controlling for updating amplified the negative direct effects of right-hemisphere stimulation, strengthening impairments after R-DLPFC (*B* = -1.365, *SE* = 0.231, *p* < .001), R-PPC (*B* = -0.964, *SE* = 0.247, *p* < .001), and R-FP stimulation (*B* = -0.954, *SE* = 0.279, *p* = .001). Indirect effects via updating were nevertheless positive for R-DLPFC (*B* = 0.553, 95%CI [0.218, 0.960], *p* < .001), with trend-level positive mediation for R-FP (*B* = 0.206, 95% CI [-0.013, 0.513], *p* = .067) and R-PPC (*B* = 0.173, 95%CI [-0.029, 0.452], *p* = .098) indicating that beneficial updating gains partially compensated for direct disruption of *Gf*. L-FP stimulation, on the other hand, showed a tendency toward reducing *Gf* (*B* = -0.406, *SE* = 0.280, *p* = .153) and a significant negative indirect effect on *Gf* via updating (*B* = -0.457, 95%CI [-0.852, -0.137], *p* = .001). After accounting for updating gains, the effect of L-FP on *Gf* was essentially null (*B* = 0.048, *SE* = 0.255, *p* = .851), indicating that the reduction in *Gf* was mediated by updating improvements.

Effects on *Gc* were more nuanced - under R-DLPFC, a significant positive indirect effect via updating (*B* = 0.455, 95% CI [0.145, 0.852], *p* = .001) was opposed by a negative but nonsignificant direct effect (*B* = -0.366, *SE* = 0.284, *p* = .203), producing a near-zero total effect (*B* = 0.090, *SE* = 0.285, *p* = .753). In contrast, R-PPC showed a trend-level positive indirect effect (*B* = 0.227, 95% CI [-0.039, 0.562], *p* = .095), alongside with a significant negative direct effect (*B* = -0.697, *SE* = 0.249, *p* = .007), resulting in trend-level negative total effect (*B* = - 0.471, *SE* = 0.277, *p* = .096).

For *Gs*, a similar pattern of offsetting effects was found. Under L-PPC stimulation, positive indirect effects via updating (*B* = 0.248, 95% CI [0.014, 0.572], *p* = .034) were canceled out by negative albeit nonsignificant direct effects (*B* = -0.368, *SE* = 0.297, *p* = .221). Likewise, under R-DLPFC, a trend-level positive indirect effect (*B* = 0.223, 95% CI [-0.015, 0.599], *p* = .069) was offset by a nonsignificant negative direct effect (*B* = -0.157, *SE* = 0.309, *p* = .614). In both instances, opposing effects canceled each other out, leaving the observed total effect near zero (*B* = -0.118, *SE* = 0.285, *p* = .680; *B* = 0.098, *SE* = 0.285, *p* = .732).

Detailed results, including DCS effects on EF (*a-paths*) and cognitive abilities (*c-paths*) together with mediation analyses, are reported in Tables S7 (left hemisphere) and S8 (right hemisphere) of the Supplementary material.

## Discussion

Study 2 demonstrates that anodal tDCS can selectively modulate both EFs and higher-order cognitive abilities, but that the effects are neither uniform across domains nor symmetric across hemispheres. Updating emerged as the only EF domain reliably enhanced by tDCS, with varying degrees of facilitation observed after DLPFC, PPC, and FP protocols. Despite L-DLPFC being the most prominent target for WM/Updating modulation in tDCS literature ^37–39,59–61^, we observed the most convincing effects after R-DLPFC stimulation. This aligns with a more recent meta-analysis demonstrating the superiority of R-DLPFC stimulation in the context of updating modulation ^41^. PPC effects on updating were somewhat less consistent and more pronounced in the left hemisphere. Notably, FP stimulation enhanced updating with the magnitude of effects comparable to single-site stimulation despite each receiving half the current intensity. This suggests that network-targeted stimulation can yield similar updating gains at a lower per-site dose, likely by enhancing coordination and information processing along the frontoparietal pathway. To the best of our knowledge, this is the first study to demonstrate that simultaneous stimulation of anterior and posterior brain regions can modulate EFs.

Null group-level tDCS effects were observed for inhibition, consistent with prior meta-analytic evidence ^38^. Unlike response inhibition, which could be reliably modulated by right IFG stimulation ^40^, our findings suggest that DLPFC stimulation has a limited potential to modulate interference control, at least when assessed with Stroop-like paradigms. Similarly, shifting performance was unaffected across all stimulation sites, again aligning with meta-analytic reports of null effects ^38,39^. One potential reason for the null effects reported in literature, including the present study, may be psychometric. Specifically, core indices of inhibition and shifting are commonly operationalized as differential RT, however, these measures typically exhibit modest reliability and can therefore attenuate or obscure any genuine tDCS-induced change.

Our study shows that the impact of anodal tDCS on higher-order cognition is not uniformly beneficial, but that direction as well as the magnitude of effects are ability-dependent. The negative effects of anodal tDCS over R-DLPFC on cognitive abilities converge with those of Sellers et al. (2015), who showed that R-DLPFC stimulation led to an overall decrease in cognitive performance on WAIS-IV, primarily due to the degradation of Perceptual Reasoning Index – an amalgam measure of *Gf* and *Gv*. However, we found that negative tDCS effects are not confined to perceptual reasoning but extend to reasoning abilities more broadly. Similar to the R-DLPFC, negative tDCS effects on fluid reasoning were observed for R-PPC as well as R-FP stimulation. These findings suggest that the negative effects of tDCS cannot be exclusively attributed to DLPFC but extend to both anterior and posterior regions of the fronto-parietal network. A likely explanation of these findings is that tDCS over the frontoparietal network may disturb coordinated, flexible interactions across right prefrontal and parietal circuits on which effective reasoning relies. On the other hand, *Gc* was largely unaffected by a single-session tDCS. The only convergent pattern was a trend-level decrease following both left and right PPC stimulation. This pattern suggests that PPC stimulation may transiently hinder performance on tests tapping *Gc*, plausibly by disrupting controlled PPC-mediated retrieval processes that support efficient access to crystallized knowledge. Although suggestive, given that both negative effects were observed only at the trend-level, these results should be interpreted with caution and require replication. In contrast to fluid reasoning, anodal tDCS over the R-DLPFC and, marginally R-PPC, were associated with better visual processing. This pattern aligns with right-lateralized domain-specific visuospatial processing, with R-DLPFC and R-PPC contributing to keeping task-relevant visual information active in WM and biasing attention toward the relevant visuospatial features of the stimuli. Finally, no reliable tDCS effects were observed for *Gs*, suggesting that tDCS has a limited potential in modulating processing speed by targeting the frontoparietal functional network.

Taken together, the results indicate that a single session of anodal tDCS applied over the frontoparietal network can modulate both EF and higher-order cognitive performance. The findings in the mediation models (discussed below) provide evidence that tDCS-induced modulation of EF, updating in particular, indirectly affects more complex cognitive functions, supporting its contributions to higher-order cognition within the frontoparietal network underlying both domains.

## General discussion

Here we presented the first comprehensive integration of psychometric and neuromodulatory approaches to examine the role of EFs in higher-order cognition. By combining psychometric modeling of the relationship between EFs and broad cognitive abilities with a systematic investigation of anodal tDCS effects across homologous brain sites, we provide convergent behavioral and mechanistic evidence on how frontoparietal control systems shape complex cognition. Study 1 highlighted the psychometric architecture of the prominent models of EFs ^13,15,58^ and cognitive abilities ^1^, offering a comprehensive mapping of the relation between the two cognitive domains. Building upon that framework, in Study 2 we presented an extensive investigation of how tDCS over key frontoparietal hubs simultaneously modulates both EFs and higher-order cognition.

Psychometric analysis showed that at the latent level, the Common EF factor reflecting the capacity to actively maintain task goals amid distraction is of particular importance for both visual processing and processing speed. Fluid reasoning, on the other hand, proved to be the only broad cognitive ability factor to show significant links with both the Common EF and Updating-specific factor, suggesting that effective reasoning relies not only on general attentional control but also on processes uniquely tied to goal maintenance and monitoring. Contrary to some earlier findings ^19,20^, crystalized ability was unrelated to any EF in particular and only weakly associated with Common EF factor, indicating that *Gc* relies more heavily than other broad cognitive abilities on domain-specific rather than domain-general processes.

The observed tDCS effects both build on and extend these psychometric findings. Specifically, the facilitatory effects for *Gv* after right-hemisphere stimulation (R-DLPFC and marginally R-PPC) were concurrently observed with enhanced updating. Crucially, when gains in updating were partialled out, *Gv* enhancement vanished, indicating a mechanistic link whereby tDCS-induced improvement in updating mediated benefits in visuospatial processing. This aligns with psychometric evidence relating *Gv* and updating, supporting the view that efficient visuospatial processing depends critically, but not exclusively, on the domain-general processes underlying maintenance and monitoring of goal-relevant representations in WM. By contrast, *Gf* performance was disrupted following R-DLPFC, R-PPC, and R-FP stimulation, concurrent with significant improvements in updating. The fact that the stimulation of the same right frontoparietal sites enhanced updating yet impaired *Gf* suggests a more complex relationship between these constructs than is often assumed based on psychometric accounts alone. Unlike *Gv*, improved updating did not translate into better *Gf* performance; instead, once tDCS-induced changes in updating were controlled, the negative effect of stimulation on *Gf* became more prominent, implying that updating gains only partly buffered and compensated for the detrimental effects on fluid reasoning. These findings extend psychometric evidence linking goal maintenance to fluid reasoning, and supports proposals that effective reasoning requires not only maintaining and monitoring representations in WM, but also other domain-general executive processes such as those supporting disengagement from no longer-relevant information^11^. Thus, it seems plausible that boosting one domain-general executive process may come at the expense of others.

Taken together, these findings suggest that updating may play distinct roles in fluid reasoning and visuospatial abilities. For instance, solving a visual puzzle requires maintaining a task-related goal in mind, in this case, an image of a missing piece, mentally manipulating it, and linearly comparing it with potential solutions. In contrast, solving a matrix or analogy problem requires not only maintaining goal-relevant information in WM but also generating, testing, and flexibly switching between competing hypotheses, as well as abandoning faulty ones. These processes likely extend beyond updating, engaging a broader executive network that controls and regulates the maintenance-disengagement dynamic, which seems central to fluid, but not visuospatial abilities. Hence, enhancing updating may push the system toward over-stability, a state optimized for information maintenance, which is well-suited to *Gv*, but less compatible with the flexibility that *Gf* requires. Still, the absence of significant tDCS effects on shifting or inhibition limits a full examination of this distinction, leaving the interpretation somewhat inconclusive and speculative. However, a general tendency for shifting disruption found both after R-DLPFC and R-PPC stimulation seems to align with this possibility. Nevertheless, the overall pattern indicates that enhancing the capacity to maintain and monitor WM representations alone is sufficient to facilitate *Gv*, yet insufficient to drive *Gf* improvement on its own. This pattern underscores the psychometric evidence towards a unique cognitive architecture of *Gf*, which appears to rely more than other abilities on a broader array of domain-general processes as well as a flexible balance between them.

Left-hemisphere stimulation did enhance updating; however, updating acted as a suppressor, masking an underlying negative impact of tDCS on *Gv*. Stimulating task-irrelevant neural pathways likely diverted cognitive resources away from those essential for visuospatial processing, with this impairment partly compensated by domain-general cognitive support arising from increased updating capacity. When those compensatory influences were partialled out, the detrimental impact of left-hemisphere stimulation on *Gv* became apparent. This pattern seems consistent with domain-specific interference, where anodal stimulation may have biased the system toward left-lateralized verbal processing, which is suboptimal for the visuospatial demands of *Gv* tasks. In general, these results illustrate competitive dynamics where tDCS enhances some domain-general executive processes, such as updating, while simultaneously diverting cognitive resources from relevant domain-specific processes, such as those underlying visuospatial information processing.

Performance tapping crystallized abilities remained largely unaffected by tDCS, both directly and indirectly, consistent with its weak psychometric relationships to EFs. Still, a similar disruption-compensation pattern appeared for R-PPC. Namely, once updating gains were partialled out, a clear disruption in *Gc* emerged, suggesting that stimulating function-irrelevant pathways can impair relevant domain-specific processes – a disruption which can be partly compensated by domain-general gains in updating. This effect plausibly reflects a transient interference with parietal mechanisms supporting controlled access to verbal representations in long-term memory.

Processing speed showed no reliable tDCS-induced changes across stimulation sites. However, when tDCS improved updating, particularly in the case of L-PPC and, at the trend level, R-DLPFC, it indirectly tended to improve *Gs* too. These findings are consistent with the notion that better updating capacity can facilitate efficiency and fluency in processing relatively simple information; yet, these indirect benefits appear to be canceled out by opposing direct modulatory influences, yielding a near-zero overall change in processing speed.

Finally, our results are consistent with the notion that group factors of cognitive abilities function as reflective outcomes of overlapping domain-general and domain-specific processes ^9^. tDCS appears to modulate these underlying processes, and observed ability-level changes arise to the extent that the modulated processes are those that a given factor weights most strongly.

When influences on domain-general and domain-specific processes align, their effects aggregate at the ability level; when they oppose, their influences on higher-order cognition can be antagonistic and cancel out, producing near-zero net behavioral change, or can mask genuine change via partial compensation. On a more general note, findings suggest that tDCS effects on complex cognition can be viewed as the net consequence of modulating multiple lower-level processes. Because these modulatory shifts in lower-level processes can differ in both direction and magnitude, their joint expression at the higher-order level may yield improvements, impairments, or near-zero change. This variability in modulatory effects on lower-level processes offers a parsimonious account of the heterogeneity and inconsistency in reported tDCS effects on higher cognition across studies.

Taken together, these findings advance both psychometric and neuromodulatory accounts of human cognition. Psychometrically, they highlight the heterogeneous architecture linking EFs to broad cognitive abilities, with *Gf* supported by both Updating-specific and general executive processes, and *Gv* and *Gs* strongly reliant solely on Common EF. Neuromodulatorily, they show that anodal tDCS over the frontoparietal network can modulate not only EFs but also affect higher-order cognition, primarily through the modulation of updating. However, these mediated benefits were frequently counteracted by direct negative effects on higher-order cognitive abilities, underscoring the complex and sometimes paradoxical consequences of tDCS on cognitive functions. Integratively, the convergence of psychometric and neuromodulatory evidence supports a hierarchical organization of cognition and a dynamic interplay between EFs and cognitive abilities at both behavioral and neurophysiological levels. Together, they present a compelling account towards EFs as a fundamental building blocks of higher cognition with updating emerging as a principal pathway through which neuromodulation shapes both facilitatory and detrimental behavioral outcomes.

## Conclusions

This study integrates psychometric and neuromodulatory approaches to elucidate the critical yet complex role of EFs in higher-order cognition. We provide evidence on closely intertwined hierarchical architecture of EFs and cognitive abilities, as well as their concurrent dependence on frontoparietal network. Updating emerged as a key domain-general mechanism underlying tDCS-induced changes in higher-order cognition. However, enhancing a single executive process does not necessarily translate to broad cognitive gain and may even hinder performance in domains that depend on a balanced interplay of different cognitive processes. These findings emphasize the need to consider both the structural organization and functional interactions of executive processes in supporting higher-order cognition and demonstrate the value of integrating psychometric and neuromodulatory approaches to advance mechanistic understanding of cognitive architecture.

## Acknowledgement.

The authors would like to thank Goran Opačić, Slađan Milanović, Marija Čolić, and Uroš Konstantinović for their help in conducting this study.

## Funding

This work was supported by EU-funded HORIZON Collaboration and Support Action TWINNIBS (101059369). MŽ and JB receive institutional support from the Ministry of Science, Technological Development and Innovation of the Republic of Serbia (451-03-137/2025-03/200163; 451-03-136/2025-03/200015). The funding body had no role in the study design, analysis and interpretation of data, writing of the report, and decision to submit the article for publication.

## Declaration of Competing interest

Authors declare that they have no competing financial or non-financial interests related to this work.

## Footnotes

1 To achieve model identification, residual variances of the verbal 3-back and letter-number tasks were constrained to small positive values.

## Notes

### Competing Interest Statement

The authors have declared no competing interest.

## References

1. McGrew, K. S. CHC theory and the human cognitive abilities project: Standing on the shoulders of the giants of psychometric intelligence research. Intelligence 37, 1–10 (2009).

2. Carroll, J. B. Human Cognitive Abilities: A Survey of Factor-Analytic Studies. (Cambridge University Press, 1993). doi:10.1017/CBO9780511571312.

3. Horn, J. L. & Blankson, N. Foundations for better understanding of cognitive abilities. in Contemporary intellectual assessment: Theories, tests, and issues (2nd edition) (eds Flanagan, D. P. & Harrison, P. L.) 41–68 (Guilford, New York, 2005).

4. Horn, J. L. & Cattell, R. B. Refinement and test of the theory of fluid and crystallized general intelligences. J. Educ. Psychol. 57, 253–270 (1966).

5. Spearman, C. ‘General Intelligence,’ Objectively Determined and Measured. Am. J. Psychol. 15, 201 (1904).

6. Thurstone, L. L. Primary Mental Abilities. (University of Chicago Press, Chicago, 1938).

7. Vernon, P. E. The Structure of Human Abilities. (Methuen, London, 1971).

8. Diamond, A. Executive Functions. Annu. Rev. Psychol. 64, 135–168 (2013).

9. Kovacs, K. & Conway, A. R. A. Process Overlap Theory: A Unified Account of the General Factor of Intelligence. Psychol. Inq. 27, 151–177 (2016).

10. Burgoyne, A. P., Mashburn, C. A., Tsukahara, J. S. & Engle, R. W. Attention control and process overlap theory: Searching for cognitive processes underpinning the positive manifold. Intelligence 91, 101629 (2022).

11. Burgoyne, A. P. & Engle, R. W. Attention Control: A Cornerstone of Higher-Order Cognition. Curr. Dir. Psychol. Sci. 29, 624–630 (2020).

12. Engle, R. W. Working Memory Capacity as Executive Attention. Curr. Dir. Psychol. Sci. 11, 19–23 (2002).

13. Friedman, N. P. & Miyake, A. Unity and diversity of executive functions: Individual differences as a window on cognitive structure. Cortex 86, 186–204 (2017).

14. Miyake, A. et al. The Unity and Diversity of Executive Functions and Their Contributions to Complex “Frontal Lobe” Tasks: A Latent Variable Analysis. Cognit. Psychol. 41, 49–100 (2000).

15. Miyake, A. & Friedman, N. P. The Nature and Organization of Individual Differences in Executive Functions: Four General Conclusions. Curr. Dir. Psychol. Sci. 21, 8–14 (2012).

16. Friedman, N. P. et al. Individual differences in executive functions are almost entirely genetic in origin. J. Exp. Psychol. Gen. 137, 201–225 (2008).

17. Friedman, N. P. & Robbins, T. W. The role of prefrontal cortex in cognitive control and executive function. Neuropsychopharmacology 47, 72–89 (2022).

18. Gustavson, D. E. et al. Genetic associations between executive functions and intelligence: A combined twin and adoption study. J. Exp. Psychol. Gen. 151, 1745–1761 (2022).

19. Friedman, N. P. et al. Not All Executive Functions Are Related to Intelligence. Psychol. Sci. 17, 172– 179 (2006).

20. Salthouse, T. A., Atkinson, T. M. & Berish, D. E. Executive Functioning as a Potential Mediator of Age-Related Cognitive Decline in Normal Adults. J. Exp. Psychol. Gen. 132, 566–594 (2003).

21. Unsworth, N. et al. Exploring the Relations Among Executive Functions, Fluid Intelligence, and Personality. J. Individ. Differ. 30, 194–200 (2009).

22. Niendam, T. A. et al. Meta-analytic evidence for a superordinate cognitive control network subserving diverse executive functions. Cogn. Affect. Behav. Neurosci. 12, 241–268 (2012).

23. Uddin, L. Q. Cognitive and behavioural flexibility: neural mechanisms and clinical considerations. Nat. Rev. Neurosci. 22, 167–179 (2021).

24. Owen, A. M., McMillan, K. M., Laird, A. R. & Bullmore, E. N-back working memory paradigm: A meta-analysis of normative functional neuroimaging studies. Hum. Brain Mapp. 25, 46–59 (2005).

25. Aron, A. R., Robbins, T. W. & Poldrack, R. A. Inhibition and the right inferior frontal cortex. Trends Cogn. Sci. 8, 170–177 (2004).

26. Aron, A. R., Robbins, T. W. & Poldrack, R. A. Inhibition and the right inferior frontal cortex: one decade on. Trends Cogn. Sci. 18, 177–185 (2014).

27. Wager, T. D., Jonides, J. & Reading, S. Neuroimaging studies of shifting attention: a meta-analysis. NeuroImage 22, 1679–1693 (2004).

28. Alvarez, J. A. & Emory, E. Executive Function and the Frontal Lobes: A Meta-Analytic Review. Neuropsychol. Rev. 16, 17–42 (2006).

29. Collette, F., Hogge, M., Salmon, E. & Van Der Linden, M. Exploration of the neural substrates of executive functioning by functional neuroimaging. Neuroscience 139, 209–221 (2006).

30. Wager, T. D. et al. Common and unique components of response inhibition revealed by fMRI. NeuroImage 27, 323–340 (2005).

31. Wager, T. D. & Smith, E. E. Neuroimaging studies of working memory: A meta-analysis. Cogn. Affect. Behav. Neurosci. 3, 255–274 (2003).

32. Jung, R. E. & Haier, R. J. The Parieto-Frontal Integration Theory (P-FIT) of intelligence: Converging neuroimaging evidence. Behav. Brain Sci. 30, 135–154 (2007).

33. Nitsche, M. A. et al. Transcranial direct current stimulation: State of the art 2008. Brain Stimulat. 1, 206–223 (2008).

34. Stagg, C. J. & Nitsche, M. A. Physiological Basis of Transcranial Direct Current Stimulation. The Neuroscientist 17, 37–53 (2011).

35. Antal, A. et al. Low intensity transcranial electric stimulation: Safety, ethical, legal regulatory and application guidelines. Clin. Neurophysiol. 128, 1774–1809 (2017).

36. Bjekić, J. et al. The subjective experience of transcranial electrical stimulation: a within-subject comparison of tolerability and side effects between tDCS, tACS, and otDCS. Front. Hum. Neurosci. 18, 1468538 (2024).

37. Hill, A. T., Fitzgerald, P. B. & Hoy, K. E. Effects of Anodal Transcranial Direct Current Stimulation on Working Memory: A Systematic Review and Meta-Analysis of Findings From Healthy and Neuropsychiatric Populations. Brain Stimulat. 9, 197–208 (2016).

38. Imburgio, M. J. & Orr, J. M. Effects of prefrontal tDCS on executive function: Methodological considerations revealed by meta-analysis. Neuropsychologia 117, 156–166 (2018).

39. De Boer, N. S. et al. The effect of non-invasive brain stimulation on executive functioning in healthy controls: A systematic review and meta-analysis. Neurosci. Biobehav. Rev. 125, 122–147 (2021).

40. Schroeder, P. A., Schwippel, T., Wolz, I. & Svaldi, J. Meta-analysis of the effects of transcranial direct current stimulation on inhibitory control. Brain Stimulat. 13, 1159–1167 (2020).

41. Narmashiri, A. & Akbari, F. The Effects of Transcranial Direct Current Stimulation (tDCS) on the Cognitive Functions: A Systematic Review and Meta-analysis. Neuropsychol. Rev. 35, 126–152 (2025).

42. Sellers, K. K. et al. Transcranial direct current stimulation (tDCS) of frontal cortex decreases performance on the WAIS-IV intelligence test. Behav. Brain Res. 290, 32–44 (2015).

43. Arif, Y., Spooner, R. K., HeinrichsLGraham, E. & Wilson, T. W. HighLdefinition transcranial direct current stimulation modulates performance and alpha/beta parietoLfrontal connectivity serving fluid intelligence. J. Physiol. 599, 5451–5463 (2021).

44. Ackerman, P. L., Beier, M. E. & Boyle, M. O. Working Memory and Intelligence: The Same or Different Constructs? Psychol. Bull. 131, 30–60 (2005).

45. Carpenter, P. A., Just, M. A. & Shell, P. What one intelligence test measures: a theoretical account of the processing in the Raven Progressive Matrices Test. Psychol. Rev. 97, 404–431 (1990).

46. Engle, R. W., Tuholski, S. W., Laughlin, J. E. & Conway, A. R. A. Working memory, short-term memory, and general fluid intelligence: A latent-variable approach. J. Exp. Psychol. Gen. 128, 309– 331 (1999).

47. Kyllonen, P. C. & Christal, R. E. Reasoning ability is (little more than) working-memory capacity?! Intelligence 14, 389–433 (1990).

48. Živanović, M., et al. The effects of offline and online prefrontal vs parietal transcranial direct current stimulation (tDCS) on verbal and spatial working memory. Neurobiol. Learn. Mem. 179, 107398 (2021).

49. Stroop, J. R. Studies of interference in serial verbal reactions. J. Exp. Psychol. 18, 643–662 (1935).

50. Purić, D. Odnos egzekutivnih funkcija i crta ličnosti. (Univerzitet u Beogradu, Filozofski fakultet, Beograd, 2013).

51. Navon, D. Forest before trees: The precedence of global features in visual perception. Cognit. Psychol. 9, 353–383 (1977).

52. Živanović, M. Efekti transkranijalne neuromodulacije fronto-parijetalne funkcionalne mreže na više kognitivne funkcije. (Univerzitet u Beogradu, Filozofski fakultet, Beograd, 2019).

53. Živanović, M., Bjekić, J. & Opačić, G. Multiple solutions test part I: Development and psychometric evaluation. Psihologija 51, 351–375 (2018).

54. Živanović, M., Bjekić, J. & Opačić, G. Multiple solutions test - part II: Evidence on construct and predictive validity. Psihologija 51, 377–396 (2018).

55. Raven, J. C., Court, J. H. & Raven, J. Manual for Raven’s Progressive Matrices and Vocabulary Scales. (H. K. Lewis Co, London, 1979).

56. Wolf, B., Momirović, K. & Džamonja, Z. KOG 3 – Baterija Testova Inteligencije. (Centar za primenjenu psihologiju, Beograd, 1992).

57. Hu, L. & Bentler, P. M. Cutoff criteria for fit indexes in covariance structure analysis: Conventional criteria versus new alternatives. Struct. Equ. Model. Multidiscip. J. 6, 1–55 (1999).

58. Miyake, A. et al. The Unity and Diversity of Executive Functions and Their Contributions to Complex “Frontal Lobe” Tasks: A Latent Variable Analysis. Cognit. Psychol. 41, 49–100 (2000).

59. Brunoni, A. R. & Vanderhasselt, M.-A. Working memory improvement with non-invasive brain stimulation of the dorsolateral prefrontal cortex: A systematic review and meta-analysis. Brain Cogn. 86, 1–9 (2014).

60. Mancuso, L. E., Ilieva, I. P., Hamilton, R. H. & Farah, M. J. Does Transcranial Direct Current Stimulation Improve Healthy Working Memory?: A Meta-analytic Review. J. Cogn. Neurosci. 28, 1063–1089 (2016).

61. Wischnewski, M., Mantell, K. E. & Opitz, A. Identifying regions in prefrontal cortex related to working memory improvement: A novel meta-analytic method using electric field modeling. Neurosci. Biobehav. Rev. 130, 147–161 (2021).

