## Supplement for "Mechanistic Link between Executive Functions and Higher Cognition: Evidence from Psychometric Modeling and Brain Stimulation"

**Supplementary material**

Table S1

*Descriptive statistics and correlations between EF measures*

| measures | *M* | *SD* | Spatial  3-back | Verbal  Stroop | Spatial  Stroop | Letter-  number | Local-  global |
| --- | --- | --- | --- | --- | --- | --- | --- |
| Verbal 3-back | 59.97 | 0.18 | .636^***^ | .201^**^ | .191^**^ | .093 | .236^***^ |
| Spatial 3-back | 55.64 | 0.20 |  | .223^**^ | .208^**^ | .045 | .169^*^ |
| Verbal Stroop | -152.52 | 124.39 |  |  | .149^*^ | .161^*^ | .253^***^ |
| Spatial Stroop | -46.68 | 39.18 |  |  |  | .022 | .067 |
| Letter-number | -317.72 | 115.83 |  |  |  |  | .272^***^ |
| Local-global | -453.64 | 175.99 |  |  |  |  |  |

*Note.* ^*^ *p* < .05; ^**^ *p* < .01; ^***^ *p* < .001

Table S2

*Descriptive statistics and correlations between cognitive ability measures*

| measures | *M* | *SD* | MTRX | FAL | RPM | S-A | CA | GSN | MSC | PZL | IT2 | SYM | VS | IT1 |
| --- | --- | --- | --- | --- | --- | --- | --- | --- | --- | --- | --- | --- | --- | --- |
| MTRX | 9.46 | 2.43 | .572 | .410^***^ | .509^***^ | .281^***^ | .316^***^ | .137^*^ | .522^***^ | .487^***^ | .469^***^ | .285^***^ | .160^*^ | .226^**^ |
| FAL | 18.90 | 4.85 |  | .770 | .464^***^ | .363^***^ | .472^***^ | .328^***^ | .439^***^ | .439^***^ | .480^***^ | .313^***^ | .230^**^ | .197^**^ |
| RPM | 14.32 | 2.38 |  |  | .654 | .193^**^ | .294^***^ | .079 | .491^***^ | .503^***^ | .507^***^ | .286^***^ | .149^*^ | .418^***^ |
| S-A | 50.09 | 4.49 |  |  |  | .850 | .329^***^ | .478^***^ | .284^***^ | .315^***^ | .201^**^ | .301^***^ | .211^**^ | .209^**^ |
| CA | 18.00 | 5.11 |  |  |  |  | .768 | .340^***^ | .307^***^ | .349^***^ | .377^***^ | .264^***^ | .046 | .185^**^ |
| GSN | 31.04 | 4.18 |  |  |  |  |  | .789 | .143^*^ | .112† | .150^*^ | .162^*^ | .179^**^ | .050 |
| MSC | 15.68 | 6.96 |  |  |  |  |  |  | .886 | .645^***^ | .609^***^ | .428^***^ | .259^***^ | .293^***^ |
| PZL | 20.27 | 6.04 |  |  |  |  |  |  |  | .827 | .626^***^ | .403^***^ | .224^**^ | .432^***^ |
| IT2 | 24.90 | 6.21 |  |  |  |  |  |  |  |  | .832 | .262^***^ | .144^*^ | .295^***^ |
| SYM | 34.79 | 8.15 |  |  |  |  |  |  |  |  |  | / | .321^***^ | .260^***^ |
| VS | 37.87 | 6.40 |  |  |  |  |  |  |  |  |  |  | / | .057 |
| IT1 | 34.38 | 3.55 |  |  |  |  |  |  |  |  |  |  |  | .724 |

*Note.* *MTRX* – Matrix test; *FAL* – Fluid analogies; *RPM* – Raven’s progressive matrices - short form; *S-A* – Synonym-antonym; *CA* – Crystalized associations; *GSN* – Lexical knowledge test; *MSC* – Mosaic; *PZL* – Puzzles; *IT2* – Spatial ability test; *SYM* – Symbol; *VS* – Visual search; *IT1* – identical figures; ^†^ *p* < .10; ^*^ *p* < .05; ^**^ *p* < .01; ^***^ *p* < .001; Diagonal values – Cronbach alpha coefficients

Table S3

*Correlations between measures of cognitive abilities and EF*

| measures | Verbal 3 back | Spatial 3-back | Verbal Stroop | Spatial Stroop | Letter-number | Local-global |
| --- | --- | --- | --- | --- | --- | --- |
| MTRX | .233^**^ | .231^**^ | .212^**^ | .124^†^ | .028 | .120^†^ |
| FAL | .181^**^ | .160^*^ | .115^†^ | -.049 | -.100 | .064 |
| RPM | .231^**^ | .325^***^ | .173^*^ | .043 | -.098 | .057 |
| S-A | .018 | .073 | .167^*^ | .064 | .015 | .153^*^ |
| CA | .125^†^ | .091 | .111 | .057 | -.112^†^ | -.010 |
| GSN | .091 | -.052 | .086 | -.043 | -.089 | .072 |
| MSC | .153^*^ | .184^**^ | .324^***^ | .090 | .112^†^ | .131^†^ |
| PZL | .233^**^ | .253^***^ | .227^**^ | .040 | .131^†^ | .168^*^ |
| IT2 | .192^**^ | .293^***^ | .260^***^ | .010 | -.018 | .083 |
| SYM | .082 | .106 | .152^*^ | .084 | .159^*^ | .163^*^ |
| VS | .220^**^ | .221^**^ | .230^**^ | .034 | .137^*^ | .177^**^ |
| IT1 | .168^*^ | .278^***^ | .209^**^ | .008 | -.084 | .165^*^ |

*Note.* *MTRX* – Matrix test; *FAL* – Fluid analogies; *RPM* – Raven’s progressive matrices - short form; *S-A* – Synonym-antonym; *CA* – Crystalized associations; *GSN* – Lexical knowledge test; *MSC* – Mosaic; *PZL* – Puzzles; *IT2* – Spatial ability test; *SYM* – Symbol; *VS* – Visual search; *IT1* – identical figures; ^†^ *p* < .10; ^*^ *p* < .05; ^**^ *p* < .01; ^***^ *p* < .001

Table S4

*Indices of reliability and equivalence for test forms and each outcome measure*

| measures | Reliability indices | | | | *ICC* |
| --- | --- | --- | --- | --- | --- |
|  | F1 | F2 | F3 | F4 |  |
| MTRX | .630^a^ | .615^a^ | .582^a^ | .629^a^ | .850^***^ |
| FAL | .821^a^ | .811^a^ | .811^a^ | .816^a^ | .925^***^ |
| S-A | .920^a^ | .916^a^ | .922^a^ | .929^a^ | .954^***^ |
| CA | .700^a^ | .720^a^ | .699^a^ | .674^a^ | .933^***^ |
| MSC | .913^a^ | .925^a^ | .904^a^ | .922^a^ | .927^***^ |
| PZL | .901^a^ | .914^a^ | .894^a^ | .912^a^ | .940^***^ |
| SYM | .629^b^ | .713^b^ | .764^b^ | .772^b^ | .924^***^ |
| VS | .736^b^ | .686^b^ | .607^b^ | .620^b^ | .892^***^ |
| Verbal 3-back | .727^b^ | .643^b^ | .597^b^ | .556^b^ | .895^***^ |
| Spatial 3-back | .682^b^ | .651^b^ | .580^b^ | .639^b^ | .901^***^ |
| Verbal Stroop | .420^b^ | .439^b^ | .270^b^ | .416^b^ | .783^***^ |
| Spatial Stroop | .211^b^ | .316^b^ | .432^b^ | .528^b^ | .763^***^ |
| Letter-number | .456^b^ | .506^b^ | .598^b^ | .584^b^ | .813^***^ |
| Local-global | .325^b^ | .462^b^ | .557^b^ | .614^b^ | .813^***^ |

*Note.* *MTRX* – Matrix test; *FAL* – Fluid analogies; *S-A* – Synonym-antonym; *CA* – Crystalized associations; *MSC* – Mosaic; *PZL* – Puzzles; *SYM* – Symbol; *VS* – Visual search; *ICC* – intraclass correlation coefficients; ^a^ Cronbach alpha; ^b^ Proportion of variance in test form accounted for by the remaining test forms (*R^2^*); ^***^ *p* < .001

Table S5

*Descriptive statistics for the outcome measures by stimulation conditions*

| hemisphere | measures | DLPFC | | PPC | | FP | | Sham | |
| --- | --- | --- | --- | --- | --- | --- | --- | --- | --- |
|  |  | *M* | *SD* | *M* | *SD* | *M* | *SD* | *M* | *SD* |
| Left | Verbal 3-back | 0.80 | 0.14 | 0.79 | 0.18 | 0.77 | 0.16 | 0.76 | 0.17 |
|  | Spatial 3-back | 0.75 | 0.21 | 0.78 | 0.24 | 0.79 | 0.17 | 0.72 | 0.22 |
|  | Verbal Stroop | -131.01 | 95.90 | -135.84 | 119.79 | -127.17 | 99.93 | -135.09 | 99.40 |
|  | Spatial Stroop | -53.20 | 44.85 | -50.62 | 47.14 | -34.63 | 40.34 | -38.27 | 43.71 |
|  | Letter-number | -269.11 | 122.24 | -249.50 | 89.53 | -248.73 | 78.85 | -263.28 | 93.96 |
|  | Local-global | -362.71 | 136.97 | -355.44 | 144.07 | -374.84 | 149.16 | -382.39 | 192.93 |
|  | MTRX | 0.69 | 0.13 | 0.67 | 0.13 | 0.67 | 0.12 | 0.67 | 0.12 |
|  | FAL | 0.68 | 0.16 | 0.68 | 0.16 | 0.67 | 0.17 | 0.70 | 0.15 |
|  | S-A | 0.69 | 0.15 | 0.69 | 0.15 | 0.70 | 0.14 | 0.68 | 0.14 |
|  | CA | 0.63 | 0.15 | 0.62 | 0.13 | 0.63 | 0.14 | 0.64 | 0.13 |
|  | MSC | 0.69 | 0.19 | 0.70 | 0.21 | 0.67 | 0.19 | 0.71 | 0.19 |
|  | PZL | 0.65 | 0.18 | 0.68 | 0.18 | 0.68 | 0.18 | 0.68 | 0.17 |
|  | SYM | 0.39 | 0.07 | 0.37 | 0.07 | 0.38 | 0.06 | 0.37 | 0.07 |
|  | VS | 0.41 | 0.07 | 0.41 | 0.08 | 0.40 | 0.06 | 0.42 | 0.07 |
| Right | Verbal 3-back | 0.82 | 0.12 | 0.80 | 0.11 | 0.78 | 0.12 | 0.76 | 0.16 |
|  | Spatial 3-back | 0.85 | 0.12 | 0.81 | 0.13 | 0.84 | 0.11 | 0.79 | 0.15 |
|  | Verbal Stroop | -112.89 | 92.28 | -139.88 | 112.37 | -140.99 | 111.52 | -124.01 | 73.77 |
|  | Spatial Stroop | -50.15 | 51.13 | -43.61 | 35.47 | -48.29 | 43.91 | -60.60 | 39.22 |
|  | Letter-number | -263.24 | 116.16 | -263.26 | 98.63 | -252.01 | 90.58 | -238.59 | 78.12 |
|  | Local-global | -350.96 | 153.38 | -357.29 | 133.16 | -332.57 | 190.68 | -337.94 | 144.59 |
|  | MTRX | 0.64 | 0.13 | 0.67 | 0.13 | 0.67 | 0.13 | 0.71 | 0.15 |
|  | FAL | 0.67 | 0.14 | 0.67 | 0.15 | 0.65 | 0.13 | 0.69 | 0.15 |
|  | S-A | 0.69 | 0.09 | 0.68 | 0.11 | 0.69 | 0.13 | 0.68 | 0.11 |
|  | CA | 0.61 | 0.15 | 0.60 | 0.14 | 0.61 | 0.13 | 0.62 | 0.14 |
|  | MSC | 0.72 | 0.18 | 0.70 | 0.19 | 0.68 | 0.20 | 0.69 | 0.18 |
|  | PZL | 0.69 | 0.15 | 0.69 | 0.16 | 0.67 | 0.20 | 0.65 | 0.19 |
|  | SYM | 0.40 | 0.07 | 0.40 | 0.06 | 0.40 | 0.08 | 0.40 | 0.06 |
|  | VS | 0.43 | 0.07 | 0.43 | 0.06 | 0.42 | 0.05 | 0.42 | 0.07 |

*Note. MTRX* – Matrix test; *FAL* – Fluid analogies; *S-A* – Synonym-antonym; *CA* – Crystalized associations; *MSC* – Mosaic; *PZL* – Puzzles; *SYM* – Symbol; *VS* – Visual search

Table S6

*Effects of condition, test, and condition* × *test interactions*

| hemisphere | measure | locus | condition | | | test | | | condition × test | | |
| --- | --- | --- | --- | --- | --- | --- | --- | --- | --- | --- | --- |
|  |  |  | *F*_(1,69)_ | *p* | η_p_^2^ | *F*_(1,69)_ | *p* | η_p_^2^ | *F*_(1,69)_ | *p* | η_p_^2^ |
| Left | *Updating* | DLPFC | 2.895 | .093 | .04 | 5.735 | .019 | .08 | 0.132 | .718 | .00 |
|  |  | PPC | 5.348 | .024 | .07 | 2.513 | .118 | .04 | 0.357 | .552 | .01 |
|  |  | FP | 4.674 | .034 | .06 | 0.443 | .508 | .01 | 2.766 | .101 | .04 |
|  | *Inhibition* | DLPFC | 0.146 | .704 | .00 | 37.726 | <.001 | .35 | 0.448 | .506 | .01 |
|  |  | PPC | 0.158 | .693 | .00 | 30.423 | <.001 | .31 | 0.124 | .726 | .00 |
|  |  | FP | 0.177 | .676 | .00 | 47.433 | <.001 | .41 | 0.024 | .876 | .00 |
|  | *Shifting* | DLPFC | 0.081 | .777 | .01 | 19.187 | <.001 | .22 | 0.276 | .601 | .00 |
|  |  | PPC | 0.983 | .325 | .01 | 30.027 | <.001 | .30 | 0.103 | .749 | .00 |
|  |  | FP | 0.287 | .594 | .00 | 35.297 | <.001 | .34 | 0.029 | .866 | .00 |
|  | *Gf* | DLPFC | 0.029 | .865 | .00 | 0.121 | .729 | .00 | 1.299 | .258 | .02 |
|  |  | PPC | 0.092 | .763 | .00 | 1.225 | .272 | .02 | 0.135 | .714 | .00 |
|  |  | FP | 0.571 | .453 | .01 | 0.691 | .409 | .01 | 0.687 | .410 | .01 |
|  | *Gc* | DLPFC | 0.001 | .982 | .00 | 7.080 | .010 | .09 | 0.070 | .792 | .00 |
|  |  | PPC | 0.313 | .578 | .01 | 11.339 | .001 | .14 | 0.514 | .476 | .01 |
|  |  | FP | 0.004 | .948 | .00 | 10.517 | .002 | .13 | 0.733 | .395 | .01 |
|  | *Gv* | DLPFC | 1.382 | .244 | .02 | 2.500 | .119 | .03 | 0.227 | .636 | .00 |
|  |  | PPC | 0.105 | .747 | .00 | 1.223 | .273 | .02 | 0.000 | .996 | .00 |
|  |  | FP | 1.661 | .202 | .02 | 0.285 | .595 | .00 | 0.755 | .388 | .01 |
|  | *Gs* | DLPFC | 0.612 | .437 | .01 | 7.362 | .008 | .10 | 2.334 | .131 | .03 |
|  |  | PPC | 0.072 | .789 | .00 | 12.273 | .001 | .15 | 0.427 | .516 | .01 |
|  |  | FP | 0.147 | .703 | .00 | 11.310 | .001 | .14 | 0.859 | .357 | .01 |
| Right | *Updating* | DLPFC | 8.071 | .006 | .10 | 2.530 | .116 | .04 | 0.006 | .939 | .00 |
|  |  | PPC | 2.127 | .149 | .03 | 1.277 | .263 | .02 | 0.186 | .668 | .00 |
|  |  | FP | 3.275 | .075 | .05 | 6.563 | .013 | .09 | 0.466 | .497 | .01 |
|  | *Inhibition* | DLPFC | 0.732 | .395 | .01 | 12.654 | <.001 | .15 | 0.001 | .979 | .00 |
|  |  | PPC | 0.002 | .967 | .00 | 35.236 | <.001 | .34 | 1.492 | .226 | .02 |
|  |  | FP | 0.035 | .853 | .00 | 38.520 | <.001 | .36 | 1.356 | .248 | .02 |
|  | *Shifting* | DLPFC | 0.915 | .342 | .01 | 12.728 | <.001 | .16 | 0.087 | .769 | .00 |
|  |  | PPC | 1.143 | .289 | .02 | 22.052 | <.001 | .24 | 0.017 | .898 | .00 |
|  |  | FP | 0.042 | .838 | .00 | 21.132 | <.001 | .23 | 0.230 | .633 | .00 |
|  | *Gf* | DLPFC | 4.258 | .043 | .06 | 0.055 | .816 | .00 | 0.837 | .363 | .01 |
|  |  | PPC | 1.756 | .190 | .02 | 0.208 | .650 | .00 | 0.072 | .789 | .00 |
|  |  | FP | 3.940 | .051 | .05 | 0.707 | .404 | .01 | 0.026 | .873 | .00 |
|  | *Gc* | DLPFC | 0.006 | .940 | .00 | 28.433 | <.001 | .29 | 0.316 | .576 | .01 |
|  |  | PPC | 1.205 | .276 | .02 | 29.121 | <.001 | .30 | 0.293 | .590 | .00 |
|  |  | FP | 0.016 | .900 | .00 | 19.727 | <.001 | .22 | 0.096 | .758 | .00 |
|  | *Gv* | DLPFC | 4.334 | .041 | .06 | 3.609 | .062 | .05 | 0.421 | .519 | .01 |
|  |  | PPC | 1.991 | .163 | .03 | 1.822 | .182 | .03 | 0.885 | .350 | .01 |
|  |  | FP | 0.234 | .630 | .00 | 4.927 | .030 | .07 | 1.386 | .243 | .02 |
|  | *Gs* | DLPFC | 0.034 | .855 | .00 | 4.697 | .034 | .06 | 0.163 | .687 | .00 |
|  |  | PPC | 0.000 | .984 | .00 | 6.612 | .012 | .09 | 0.433 | .513 | .01 |
|  |  | FP | 0.003 | .955 | .00 | 3.100 | .083 | .04 | 0.016 | .900 | .00 |

Table S7

*Mediation analyses for the* ***left hemisphere***

| Condition | EF | *a* | Abilities | *c* | *c’* | *b* | *a*×*b [95%CI]* |
| --- | --- | --- | --- | --- | --- | --- | --- |
| L-DLPFC | *updating* | *B* = 0.643  *SE* = 0.270  *p* = .021 | *Gf* | *B* = 0.161  *SE* = 0.285  *p* = .573 | *B* = 0.152  *SE* = 0.301  *p* = .615 | *B* = 0.014  SE = 0.152  *p* = .926 | *B* = 0.007  [-0.201, 0.220]  *p* = .934 |
|  |  |  | *Gc* | *B* = -0.020  *SE* = 0.286  *p* = .945 | *B* = -0.103  *SE* = 0.3  *p* = .734 | *B* = 0.129  *SE* = 0.151  *p* = .400 | *B* = 0.081  [-0.112, 0.324]  *p* = .398 |
|  |  |  | *Gv* | *B* = -0.353  *SE* = 0.281  *p* = .215 | *B* = -0.706  *SE* = 0.252  *p* = .007 | *B* = 0.549  *SE* = 0.128  *p* < .001 | *B* = 0.352  [0.056, 0.726]  *p* = .014 |
|  |  |  | *Gs* | *B* = 0.383  *SE* = 0.280  *p* = .178 | *B* = 0.331  *SE* = 0.295  *p* = .268 | *B* = 0.080  *SE* = 0.149  *p* = .593 | *B* = 0.050  [-0.145, 0.277]  *p* = .595 |
|  | *inhibition* | *B* = -0.393  *SE* = 0.280  *p* = .167 | *Gf* | *B* = 0.161  *SE* = 0.285  *p* = .573 | *B* = 0.105  *SE* = 0.288  *p* = .718 | *B* = -0.145  *SE* = 0.145  *p* = .324 | *B* = 0.057  [-0.072, 0.258]  *p* = .420 |
|  |  |  | *Gc* | *B* = -0.020  *SE* = 0.286  *p* = .945 | *B* = -0.011  *SE* = 0.291  *p* = .970 | *B* = 0.023  *SE* = 0.147  *p* = .877 | *B* = -0.009  [-0.172, 0.144]  *p* = .906 |
|  |  |  | *Gv* | *B* = -0.353  *SE* = 0.281  *p* = .215 | *B* = -0.189  *SE* = 0.261  *p* = .471 | *B* = 0.418  *SE* = 0.132  *p* = .003 | *B* = -0.163  [-0.460, 0.062]  *p* = .162 |
|  |  |  | *Gs* | *B* = 0.383  *SE* = 0.280  *p* = .178 | *B* = 0.430  *SE* = 0.284  *p* = .136 | *B* = 0.120  *SE* = 0.143  *p* = .406 | *B* = -0.047  [-0.240, 0.082]  *p* = .509 |
|  | *shifting* | *B* = 0.062  *SE* = 0.286  *p* = .828 | *Gf* | *B* = 0.161  *SE* = 0.285  *p* = .573 | *B* = 0.173  *SE* = 0.280  *p* = .540 | *B* = -0.184  *SE* = 0.141  *p* = .200 | *B* = -0.013  [-0.162, 0.122]  *p* = .835 |
|  |  |  | *Gc* | *B* = -0.020  *SE* = 0.286  *p* = .945 | *B* = -0.022  *SE* = 0.286  *p* = .940 | *B* = 0.029  *SE* = 0.144  *p* = .843 | *B* = 0.001  [-0.086, 0.095]  *p* = .975 |
|  |  |  | *Gv* | *B* = -0.353  *SE* = 0.281  *p* = .215 | *B* = -0.390  *SE* = 0.227  *p* = .092 | *B* = 0.582  *SE* = 0.115  *p* < .001 | *B* = 0.037  [-0.292, 0.375]  *p* = .812 |
|  |  |  | *Gs* | *B* = 0.383  *SE* = 0.280  *p* = .178 | *B* = 0.359  *SE* = 0.258  *p* = .171 | *B* = 0.381  *SE* = 0.131  *p* = .005 | *B* = 0.024  [-0.207, 0.258]  *p* = .813 |
| L-PPC | *updating* | *B* = 0.784  *SE* = 0.262  *p* = .004 | *Gf* | *B* = -0.154  *SE* = 0.285  *p* = .590 | *B* = -0.207  *SE* = 0.310  *p* = .508 | *B* = 0.067  *SE* = 0.156  *p* = .672 | *B* = 0.050  [-0.197, 0.318]  *p* = .675 |
|  |  |  | *Gc* | *B* = -0.470  *SE* = 0.278  *p* = .097 | *B* = -0.561  *SE* = 0.300  *p* = .068 | *B* = 0.117  *SE* = 0.152  *p* = .445 | *B* = 0.090  [-0.144, 0.359]  *p* = .439 |
|  |  |  | *Gv* | *B* = -0.108  *SE* = 0.285  *p* = .707 | *B* = -0.508  *SE* = 0.274  *p* = .070 | *B* = 0.510  *SE* = 0.139  *p* < .001 | *B* = 0.399  [0.108, 0.774]  *p* = .002 |
|  |  |  | *Gs* | *B* = -0.118  *SE* = 0.285  *p* = .680 | *B* = -0.368  SE = 0.297  *p* = .221 | *B* = 0.319  *SE* = 0.150  *p* = .039 | *B* = 0.248  [0.014, 0.572]  *p* = .034 |
|  | *inhibition* | *B* = -0.350  *SE* = 0.281  *p* = .219 | *Gf* | *B* = -0.154  *SE* = 0.285  *p* = .590 | *B* = -0.156  *SE* = 0.289  *p* = .592 | *B* = -0.004  *SE* = 0.146  *p* = .977 | *B* = 0.001  [-0.142, 0.149]  *p* = .979 |
|  |  |  | *Gc* | *B* = -0.470  *SE* = 0.278  *p* = .097 | *B* = -0.332  *SE* = 0.259  *p* = .205 | *B* = 0.392  *SE* = 0.131  *p* = .004 | *B* = -0.136  [-0.415, 0.077]  *p* = .214 |
|  |  |  | *Gv* | *B* = -0.108  *SE* = 0.285  *p* = .707 | *B* = -0.129  *SE* = 0.289  *p* = .658 | *B* = -0.060  *SE* = 0.146  *p* = .683 | *B* = 0.021  [-0.110, 0.186]  *p* = .759 |
|  |  |  | *Gs* | *B* = -0.118  *SE* = 0.285  *p* = .680 | *B* = -0.139  *SE* = 0.289  *p* = .633 | *B* = -0.059  *SE* = 0.146  *p* = .690 | *B* = 0.020  [-0.110, 0.185]  *p* = .767 |
|  | *shifting* | *B* = 0.410  *SE* = 0.280  *p* = .149 | *Gf* | *B* = -0.154  *SE* = 0.285  *p* = .590 | *B* = -0.294  *SE* = 0.275  *p* = .290 | *B* = 0.340  *SE* = 0.139  *p* = .018 | *B* = 0.138  [-0.044, 0.403]  *p* = .160 |
|  |  |  | *Gc* | *B* = -0.470  *SE* = 0.278  *p* = .097 | *B* = -0.539  *SE* = 0.280  *p* = .060 | *B* = 0.170  *SE* = 0.141  *p* = .235 | *B* = 0.068  [-0.052, 0.264]  *p* = .341 |
|  |  |  | *Gv* | *B* = -0.108  *SE* = 0.285  *p* = .707 | *B* = -0.352  *SE* = 0.237  *p* = .144 | *B* = 0.595  *SE* = 0.120  *p* < .001 | *B* = 0.244  [-0.079, 0.609]  *p* = .145 |
|  |  |  | *Gs* | *B* = -0.118  *SE* = 0.285  *p* = .680 | *B* = -0.237  *SE* = 0.280  *p* = .401 | *B* = 0.289  *SE* = 0.141  *p* = .046 | *B* = 0.118  [-0.040, 0.362]  *p* = .182 |
| L-FP | *updating* | *B* = 0.784  *SE* = 0.262  *p* = .004 | *Gf* | *B* = -0.406  *SE* = 0.280  *p* = .153 | *B* = 0.048  *SE* = 0.255  *p* = .851 | *B* = -0.580  *SE* = 0.129  *p* < .001 | *B* = -0.457  [-0.852 -0.137]  *p* = .001 |
|  |  |  | *Gc* | *B* = 0.028  *SE* = 0.286  *p* = .922 | *B* = 0.039  *SE* = 0.311  *p* = .900 | *B* = -0.015  *SE* = 0.157  *p* = .926 | *B* = -0.014  [-0.272, 0.244]  *p* = .918 |
|  |  |  | *Gv* | *B* = -0.418  *SE* = 0.279  *p* = .141 | *B* = -0.730  *SE* = 0.282  *p* = .013 | *B* = 0.398  *SE* = 0.143  *p* = .007 | *B* = 0.310  [0.058, 0.652]  *p* = .007 |
|  |  |  | *Gs* | *B* = -0.211  *SE* = 0.284  *p* = .461 | *B* = -0.100  *SE* = 0.307  *p* = .749 | *B* = -0.143  *SE* = 0.155  *p* = .361 | *B* = -0.115  [-0.400, 0.123]  *p* = .334 |
|  | *inhibition* | *B* = 0.224  *SE* = 0.284  *p* = .434 | *Gf* | *B* = -0.406  *SE* = 0.280  *p* = .153 | *B* = -0.396  *SE* = 0.281  *p* = .165 | *B* = -0.044  *SE* = 0.142  *p* = .757 | *B* = -0.011  [-0.133, 0.086]  *p* = .846 |
|  |  |  | *Gc* | *B* = 0.028  *SE* = 0.286  *p* = .922 | *B* = -0.004  *SE* = 0.285  *p* = .989 | *B* = 0.142  *SE* = 0.144  *p* = .329 | *B* = 0.031  [-0.074, 0.187]  *p* = .626 |
|  |  |  | *Gv* | *B* = -0.418  *SE* = 0.279  *p* = .141 | *B* = -0.409  *SE* = 0.281  *p* = .152 | *B* = -0.041  *SE* = 0.142  *p* = .773 | *B* = -0.011  [-0.132, 0.088]  *p* = .849 |
|  |  |  | *Gs* | *B* = -0.211  *SE* = 0.284  *p* = .461 | *B* = -0.232  *SE* = 0.285  *p* = .418 | *B* = 0.096  *SE* = 0.144  *p* = .507 | *B* = 0.021  [-0.074, 0.158]  *p* = .726 |
|  | *shifting* | *B* = 0.295  *SE* = 0.283  *p* = .301 | *Gf* | *B* = -0.406  *SE* = 0.280  *p* = .153 | *B* = -0.372  *SE* = 0.281  *p* = .192 | *B* = -0.117  *SE* = 0.142  *p* = .414 | *B* = -0.036  [-0.197, 0.070]  *p* = .563 |
|  |  |  | *Gc* | *B* = 0.028  *SE* = 0.286  *p* = .922 | *B* = -0.018  *SE* = 0.285  *p* = .949 | *B* = 0.157  *SE* = 0.144  *p* = .281 | *B* = 0.045  [-0.065, 0.220]  *p* = .506 |
|  |  |  | *Gv* | *B* = -0.418  *SE* = 0.279  *p* = .141 | *B* = -0.564  *SE* = 0.245  *p* = .026 | *B* = 0.493  *SE* = 0.124  *p* < .001 | *B* = 0.145  [-0.125, 0.455]  *p* = .305 |
|  |  |  | *Gs* | *B* = -0.211  *SE* = 0.284  *p* = .461 | *B* = -0.326  *SE* = 0.265  *p* = .224 | *B* = 0.390  *SE* = 0.134  *p* = .005 | *B* = 0.115  [-0.097, 0.383]  *p* = .308 |

Table S8

*Mediation analyses for the* ***right hemisphere***

| Condition | EF | *a* | Abilities | *c* | *c’* | *b* | *a*×*b [95%CI]* |
| --- | --- | --- | --- | --- | --- | --- | --- |
| R-DLPFC | *updating* | *B* = 0.888  *SE* = 0.255  *p* = .001 | *Gf* | *B* = -0.811  *SE* = 0.261  *p* = .003 | *B* = -1.365  *SE* = 0.231  *p* < .001 | *B* = 0.623  *SE* = 0.117  *p* < .001 | *B* = 0.553  [0.218, 0.960]  *p* < .001 |
|  |  |  | *Gc* | *B* = 0.090  *SE* = 0.285  *p* = .753 | *B* = -0.366  *SE* = 0.284  *p* = .203 | *B* = 0.514  *SE* = 0.143  *p* < .001 | *B* = 0.455  [0.145, 0.852]  *p* = .001 |
|  |  |  | *Gv* | *B* = 0.735  *SE* = 0.265  *p* = .008 | *B* = 0.278  *SE* = 0.258  *p* = .287 | *B* = 0.515  *SE* = 0.130  *p* < .001 | *B* = 0.456  [0.157, 0.834]  *p* < .001 |
|  |  |  | *Gs* | *B* = 0.098  *SE* = 0.285  *p* = .732 | *B* = -0.157  *SE* = 0.309  *p* = .614 | *B* = 0.287  *SE* = 0.156  *p* = .072 | *B* = 0.223  [-0.015, 0.599]  *p* = .069 |
|  | *inhibition* | *B* = 0.382  *SE* = 0.280  *p* = .179 | *Gf* | *B* = -0.811  *SE* = 0.261  *p* = .003 | *B* = -0.889  *SE* = 0.259  *p* = .001 | *B* = 0.204  *SE* = 0.131  *p* = .127 | *B* = 0.077  [-0.044, 0.272]  *p* = .283 |
|  |  |  | *Gc* | *B* = 0.090  *SE* = 0.285  *p* = .753 | *B* = -0.044  *SE* = 0.273  *p* = .873 | *B* = 0.351  *SE* = 0.138  *p* = .014 | *B* = 0.133  [-0.054, 0.400]  *p* = .190 |
|  |  |  | *Gv* | *B* = 0.735  *SE* = 0.265  *p* = .008 | *B* = 0.596  SE = 0.250  *p* = .021 | *B* = 0.363  *SE* = 0.126  *p* = .006 | *B* = 0.138  [-0.056, 0.403]  *p* = .183 |
|  |  |  | *Gs* | *B* = 0.098  *SE* = 0.285  *p* = .732 | *B* = 0.191  *SE* = 0.282  *p* = .502 | *B* = -0.243  *SE* = 0.143  *p* = .095 | *B* = -0.095  [-0.321, 0.047]  *p* = .244 |
|  | *shifting* | *B* = -0.354  *SE* = 0.281  *p* = .214 | *Gf* | *B* = -0.811  *SE* = 0.261  *p* = .003 | *B* = -0.724  *SE* = 0.255  *p* = .007 | *B* = 0.245  *SE* = 0.129  *p* = .063 | *B* = -0.086  [-0.301, 0.051]  *p* = .266 |
|  |  |  | *Gc* | *B* = 0.090  *SE* = 0.285  *p* = .753 | *B* = 0.152  *SE* = 0.286  *p* = .598 | *B* = 0.174  *SE* = 0.144  *p* = .234 | *B* = -0.061  [-0.263, 0.061]  *p* = .403 |
|  |  |  | *Gv* | *B* = 0.735  *SE* = 0.265  *p* = .008 | *B* = 0.977  *SE* = 0.185  *p* < .001 | *B* = 0.686  *SE* = 0.094  *p* < .001 | *B* = -0.241  [-0.638, 0.131]  *p* = .208 |
|  |  |  | *Gs* | *B* = 0.098  *SE* = 0.285  *p* = .732 | *B* = 0.229  *SE* = 0.270  *p* = .401 | *B* = 0.370  *SE* = 0.137  *p* = .009 | *B* = -0.130  [-0.405, 0.072]  *p* = .212 |
| R-PPC | *updating* | *B* = 0.461  *SE* = 0.278  *p* = .104 | *Gf* | *B* = -0.792  *SE* = 0.262  *p* = .004 | *B* = -0.964  *SE* = 0.247  *p* < .001 | *B* = 0.372  *SE* = 0.125  *p* = .005 | *B* = 0.173  [-0.029, 0.452]  *p* = .098 |
|  |  |  | *Gc* | *B* = -0.471  *SE* = 0.277  *p* = .096 | *B* = -0.697  *SE* = 0.249  *p* = .007 | *B* = 0.489  *SE* = 0.126  *p* < .001 | *B* = 0.227  [-0.039, 0.562]  *p* = .095 |
|  |  |  | *Gv* | *B* = 0.551  *SE* = 0.274  *p* = .050 | *B* = 0.324  *SE* = 0.245  *p* = .191 | *B* = 0.493  *SE* = 0.124  *p* < .001 | *B* = 0.229  [-0.039, 0.562]  *p* = .095 |
|  |  |  | *Gs* | *B* = -0.005  *SE* = 0.286  *p* = .986 | *B* = -0.109  *SE* = 0.287  *p* = .706 | *B* = 0.225  *SE* = 0.145  *p* = .126 | *B* = 0.105  [-0.040, 0.344]  *p* = .206 |
|  | *inhibition* | *B* = 0.351  *SE* = 0.281  *p* = .218 | *Gf* | *B* = -0.792  *SE* = 0.262  *p* = .004 | *B* = -0.886  *SE* = 0.255  *p* = .001 | *B* = 0.266  *SE* = 0.129  *p* = .044 | *B* = 0.093  [-0.052, 0.309]  *p* = .252 |
|  |  |  | *Gc* | *B* = -0.471  *SE* = 0.277  *p* = .096 | *B* = -0.474  *SE* = 0.282  *p* = .099 | *B* = 0.008  *SE* = 0.142  *p* = .953 | *B* = 0.001  [-0.127, 0.136]  *p* = .970 |
|  |  |  | *Gv* | *B* = 0.551  *SE* = 0.274  *p* = .050 | *B* = 0.367  *SE* = 0.235  *p* = .125 | *B* = 0.525  *SE* = 0.119  *p* < .001 | *B* = 0.184  [-0.100, 0.511]  *p* = .219 |
|  |  |  | *Gs* | *B* = -0.005  *SE* = 0.286  *p* = .986 | *B* = -0.075  *SE* = 0.285  *p* = .793 | *B* = 0.200  *SE* = 0.144  *p* = .171 | *B* = 0.069  [-0.054, 0.269]  *p* = .353 |
|  | *shifting* | *B* = -0.389  *SE* = 0.280  *p* = .171 | *Gf* | *B* = -0.792  *SE* = 0.262  *p* = .004 | *B* = -0.691  *SE* = 0.257  *p* = .010 | *B* = 0.260  *SE* = 0.130  *p* = .051 | *B* = -0.100  [-0.327, 0.042]  *p* = .210 |
|  |  |  | *Gc* | *B* = -0.471  *SE* = 0.277  *p* = .096 | *B* = -0.408  *SE* = 0.279  *p* = .150 | *B* = 0.162  *SE* = 0.141  *p* = .258 | *B* = -0.062  [-0.264, 0.063]  *p* = .393 |
|  |  |  | *Gv* | *B* = 0.551  *SE* = 0.274  *p* = .050 | *B* = 0.748  *SE* = 0.239  *p* = .003 | *B* = 0.507  *SE* = 0.121  *p* < .001 | *B* = -0.196  [-0.526, 0.077]  *p* = .167 |
|  |  |  | *Gs* | *B* = -0.005  *SE* = 0.286  *p* = .986 | *B* = 0.172  *SE* = 0.261  *p* = .513 | *B* = 0.454  *SE* = 0.132  *p* = .001 | *B* = -0.176  [-0.487, 0.069]  *p* = .168 |
| R-FP | *updating* | *B* = 0.790  *SE* = 0.262  *p* = .004 | *Gf* | *B* = -0.750  *SE* = 0.264  *p* = .007 | *B* = -0.954  *SE* = 0.279  *p* = .001 | *B* = 0.258  SE = 0.141  *p* = .073 | *B* = 0.206  [-0.013, 0.513]  *p* = .067 |
|  |  |  | *Gc* | *B* = 0.080  *SE* = 0.285  *p* = .780 | *B* = -0.039  *SE* = 0.308  *p* = .901 | *B* = 0.150  *SE* = 0.156  *p* = .339 | *B* = 0.121  [-0.128, 0.421]  *p* = .326 |
|  |  |  | *Gv* | *B* = 0.221  *SE* = 0.284  *p* = .440 | *B* = 0.076  *SE* = 0.305  *p* = .803 | *B* = 0.183  *SE* = 0.154  *p* = .240 | *B* = 0.147  [-0.094, 0.454]  *p* = .232 |
|  |  |  | *Gs* | *B* = -0.038  *SE* = 0.286  *p* = .894 | *B* = -0.213  *SE* = 0.305  *p* = .489 | *B* = 0.221  *SE* = 0.154  *p* = .158 | *B* = 0.177  [-0.063, 0.495]  *p* = .153 |
|  | *inhibition* | *B* = 0.153  *SE* = 0.285  *p* = .593 | *Gf* | *B* = -0.750  *SE* = 0.264  *p* = .007 | *B* = -0.686  *SE* = 0.237  *p* = .006 | *B* = -0.420  *SE* = 0.120  *p* < .001 | *B* = -0.066  [-0.328, 0.177]  *p* = .579 |
|  |  |  | *Gc* | *B* = 0.080  *SE* = 0.285  *p* = .780 | *B* = 0.131  *SE* = 0.270  *p* = .628 | *B* = -0.335  *SE* = 0.136  *p* = .018 | *B* = -0.053  [-0.279, 0.144]  *p* = .586 |
|  |  |  | *Gv* | *B* = 0.221  *SE* = 0.284  *p* = .440 | *B* = 0.213  *SE* = 0.284  *p* = .458 | *B* = 0.054  *SE* = 0.144  *p* = .709 | *B* = 0.007  [-0.083, 0.119]  *p* = .885 |
|  |  |  | *Gs* | *B* = -0.038  *SE* = 0.286  *p* = .894 | *B* = 0.021  *SE* = 0.265  *p* = .938 | *B* = -0.383  *SE* = 0.134  *p* = .006 | *B* = -0.061  [-0.309, 0.164]  *p* = .580 |
|  | *shifting* | *B* =-0.167  *SE* =0.285  *p* = .561 | *Gf* | *B* = -0.750  *SE* = 0.264  *p* = .007 | *B* = -0.765  *SE* = 0.264  *p* = .006 | *B* = -0.093  *SE* = 0.133  *p* = .491 | *B* = 0.015  [-0.076, 0.142]  *p* = .794 |
|  |  |  | *Gc* | *B* = 0.080  *SE* = 0.285  *p* = .780 | *B* = 0.115  *SE* = 0.280  *p* = .685 | *B* = 0.206  *SE* = 0.142  *p* = .152 | *B* = -0.034  [-0.214, 0.095]  *p* = .640 |
|  |  |  | *Gv* | *B* = 0.221  *SE* = 0.284  *p* = .440 | *B* = 0.288  *SE* = 0.261  *p* = .274 | *B* = 0.402  *SE* = 0.132  *p* = .004 | *B* = -0.067  [-0.331, 0.163]  *p* = .564 |
|  |  |  | *Gs* | *B* = -0.038  *SE* = 0.286  *p* = .894 | *B* = -0.002  *SE* = 0.280  *p* = .995 | *B* = 0.218  *SE* = 0.141  *p* = .130 | *B* = -0.036  [-0.221, 0.097]  *p* = .627 |
